# Enhanced *in vitro* aggregation, but not phase separation, of TDP-43 and its C-terminal fragments generate deep-blue autofluorescence

**DOI:** 10.1101/2024.11.23.624964

**Authors:** Preethi Saravanan, Vidhya Bharathi, Priyadarshini Veerabhadraswamy, Basant K Patel

## Abstract

As misfolding and aggregation of the RNA/DNA-binding protein, TDP-43, are linked to devastating TDP-43 proteinopathies like amyotrophic lateral sclerosis (ALS), distinction of the nature of the aggregated TDP-43 species being liquid-like non-pathogenic or solid-like pathogenic is important for mechanistic elucidation and therapeutic targeting. Here, we examined if *in vitro* enhancement of the TDP-43 aggregation can generate or enhance intrinsic deep-blue autofluorescence (dbAF) previously reported for a few other protein aggregates and whether dbAF is emitted by all or only liquid-like or solid-like TDP-43 aggregates. Using thioflavin-T fluorescence, turbidimetry, atomic force microscopy and fluorescence microscopy of Alexa Fluor-labelled protein, we first tested the *in vitro* enhancement of the aggregation of the full-length TDP-43 and its two C-terminal fragments (CTFs), TDP-43^2C^ (aa: 193-414) and the TDP-43-low complexity domain (LCD) (aa: 274-414). We find that presence of metal ions, Zn^2+^ or Mn^2+^, that are also linked to ALS-associated metal dyshomeostasis, or addition of a kosmotropic anion, SO_4_^2-^, enhance the *in vitro* solid-like aggregations of the full-length TDP-43 and TDP-43^2C^ that also concurrently enhance emission of dbAF. In contrast, Alexa fluor-633-labeled-TDP-43-LCD underwent a quick phase separation into globular structures in presence of Zn^2+^ ions and the phase-separated species failed to emit dbAF but upon further incubation when matured into solid-like irregular, but non-amyloid nature aggregates, it emitted dbAF. Strikingly, we find that the TDP-43 aggregates of both amyloid and non-amyloid nature, but not the oligomers or the phase-separated droplets of TDP-43, manifest dbAF. Overall, the observed *in vitro* enhancement of aggregation leading to concurrent enhancement of dbAF can enable a label-free easy detection and may facilitate distinguishing of potentially pathogenic versus non-pathogenic TDP-43 aggregates.

## 1. Introduction

Certain misfolded proteins can undergo oligomerization and form insoluble aggregates termed as amyloids that possess β-sheet-rich structural conformation [1]. The presence of intracellular or extracellular protein inclusions in the nervous system is implicated as causative agent in various neurodegenerative diseases such as Alzheimer’s disease, Parkinson’s disease and amyotrophic lateral sclerosis (ALS) [2]. TDP-43 (Transactive response DNA-binding protein), whose inclusions are implicated in ALS and several other neurodegenerative diseases, is an RNA/DNA binding protein consisting of a N-terminal domain (NTD), two RNA-recognition motifs (RRM1 and RRM2) and a C-terminal domain which is an intrinsically disordered, prion-like, low complexity domain (LCD) [3]. Notably, TDP-43 can undergo proteolytic cleavage by caspases or calpain to generate truncated forms of TDP-43 termed C-terminal fragments (CTFs) such as TDP-35 (80-414), TDP-25 (175-414) and TDP-16 (274-414) which can be found in the ALS or TDP-43-linked other neurodegenerative disease patients and even get generated in the TDP-43-expressing various model systems [4,5]. Specific conserved core regions of TDP-43 protein and its CTFs have been found to phase-separate and also form amyloid-like fibrils *in vitro* [6–11]. Metal ion dyshomeostasis and the consequent production of ROS can also contribute to the onset of several neurodegenerative diseases including ALS [12]. Recently, levels of nine metals including zinc and manganese were found to be elevated in the CSF of the ALS patients [13]. Likewise, increase in the levels of Zn^2+^, Mn^2+^, and Cu^2+^ metal ions were also observed previously in the spinal cord of mutant TDP-43^A315T^-expressing transgenic mice [14]. Previously, Zn^2+^ ion was shown to bind to recombinantly purified RRM1-2 domain fragment of TDP-43, promoting its amyloid-like *in vitro* aggregation and a binding site for Zn^2+^ was predicted in the linker region between RRM2 domain and LCD of TDP-43 [15,16]. Earlier, we also showed that an increase in the Zn^2+^ concentration accelerated the formation of *in vitro* irregular-shaped aggregates species of a C-terminal fragment of TDP-43 protein, termed TDP-43^2C^, that encompasses the RRM2 domain and LCD of TDP-43 thereby supporting that Zn^2+^ can affect the aggregation of TDP-43 fragments [17].

Intrinsic fluorescence spectrometry of the aromatic amino acids such as tryptophan and tyrosine have been traditionally utilized to study protein conformation, however, they emit fluorescence in the UV region thwarting any convenient visualization by fluorescence microscopy and thereby limiting the application [18]. Attracting significant interest, few studies have shown that *in vitro*-made aggregates of certain amyloidogenic proteins such as lysozyme, tau and Aβ peptide upon amyloid aggregation, when excited at near UV range can emit a novel deep-blue autofluorescence (dbAF) in the visible range of light that can be easily imaged by fluorescence microscopy [19–21]. Although, the possible origin of the dbAF upon aggregation remains to be vividly elucidated, certain contributing factors have been hypothesized such as the hydrogen bonds between the fibrils, carbonyl groups of proteins and oxidation of certain amino acids [23–25]. Notably, certain extrinsic dyes such as thioflavin-T (Th-T), Congo-red and thioflavin-S are commonly used for amyloid detection, however, some of these dyes can also occasionally perturb the aggregation process of certain proteins [26,27]. In this light, the possibility of employing the dbAF detection as a label-free, non-invasive detection of protein aggregates would help avoid any interference in the aggregation as observed from the extrinsic dyes and also facilitate *in vitro* screening of inhibitors or enhancers of protein aggregation without the chance of any interaction of the small molecules with the extrinsic dyes that may skew the results.

So far, a study observed that the C-terminal prion-like domain of TDP-43 with wild-type or mutant sequence, can *in vitro* emit enhanced dbAF when aggregated [22], however, whether the aggregates of full-length TDP-43 or any other CTFs of TDP-43 can also emit dbAF and furthermore, whether all types or only specific types of the protein assemblies of TDP-43, or its CTFs, cause emission of dbAF, remain to be investigated. As dbAF emission upon aggregation of TDP-43^2C^, either when alone or in the presence of aggregation enhancers, has not been reported so far, here we examined the effects of known *in vitro* aggregation enhancers such as Zn^2+^ and SO_4_^2-^ on dbAF of TDP-43^2C^ [10,28]. Furthermore, as the effects of Zn^2+^ have not yet been examined on the aggregation of full-length TDP-43 and TDP-43-LCD, we also investigated the effect of Zn^2+^ on the aggregation of these proteins using some of the widely used tools such as thioflavin-T fluorescence spectroscopy, turbidimetry, atomic force microscopy and fluorescence microscopy and then examined if their aggregations are also accompanied by emission of dbAF. Additionally, we also investigated the effect of another, yet unexplored, metal ion, Mn^2+^, the aggregation and dbAF generation by full-length TDP-43 and TDP-43^2C^. Finally, we examined if phase separation of Alexa fluor-633-labelled-TDP-43-LCD protein in the presence or absence of Zn^2+^ ions can also generate dbAF in order to examine if the dbAF emission is specific for solid-like protein aggregates that can be potentially pathogenic or can even detect phase-separated protein assembly droplets that are usually non-pathogenic and may even be physiologically functional [29,30].

## 2. Materials and methods

### 2.1. Materials

Ni-NTA agarose beads were purchased from Qiagen (USA). Guanidine hydrochloride (GuHCl) was purchased from SRL (India). Zinc chloride, dimethyl sulfoxide (DMSO), urea, yeast extract, tryptone and agarose were procured from HiMedia (India). Bradford reagent was purchased from Bio-Rad (USA). Ampicillin, chloramphenicol, phenylmethanesulfonyl fluoride (PMSF), imidazole, thioflavin-T, dithiothreitol (DTT), manganese chloride, N-lauroylsarcosine sodium salt and isopropyl β-D-1-thiogalactopyranoside (IPTG), Ethylene diamine tetra acetic acid (EDTA) disodium salt dihydrate were procured from Sigma (USA). Sodium sulfate was purchased from Finar (India). AFM mica sheets (Grade V) were purchased from SPI Supplies, USA.

### 2.2. Plasmid construction

A recombinant expression plasmid encoding TDP-43-low complexity domain (LCD) was constructed by PCR amplifying the aa: 274-414 coding region from the template plasmid, *pET15b-His-TDP-43-2C* (encoding aa: 193-414 of TDP-43 and encompassing the domains RRM2 and LCD of TDP-43) [10] using the forward primer: 5’ *TATATA<u>CTCGAG</u>GGAAGATTTGGTGGTAATCC* 3’ (with an *XhoI* restriction site) and the reverse primer: 5’*ATATTA<u>GGATCC</u>CTACATTCCCCAGCCAGAAG* 3’ (with a *BamHI* restriction site). The TDP-43-LCD sequence with the *BamHI* and *XhoI* site was sub-cloned into the multiple cloning site (*BamHI* and *XhoI*) of the *pET15b* vector with an N-terminal 6x-His tag. The obtained plasmid was named *pET15b-His-TDP-43-LCD* and used further for the recombinant protein expression.

### 2.3. Recombinant expression and purification of full-length TDP-43 and its C-terminal fragments

For the recombinant protein expression, the plasmids *pET15b-His-TDP-43* (encoding full-length TDP-43), *pET15b-His-TDP-43-2C* (encoding aa: 193-414 of TDP-43 encompassing the RRM2 and LCD of TDP-43) (kind gifts from Prof. Yoshiaki Furukawa, Keio University, Japan) and *pET15b-His-TDP-43-LCD* (encoding the low complexity domain (LCD) (aa: 274-414) of TDP-43) were transformed into competent *Rosetta 2 (DE3) E. coli* cells (Novagen, USA). Recombinant full-length TDP-43, TDP-43^2C^ protein and TDP-43-LCD were expressed and purified under denaturing conditions using nickel affinity chromatography as reported earlier [10,28,31]. Briefly, protein expression was carried out by induction with 1 mM IPTG for 4h. Then, the protein-expressing cells were harvested and lysed by ultra-sonication in the presence of the lysis buffer (6M GuHCl and 1 mM PMSF, pH 7.5). The cell lysate was pre-cleared and loaded onto a pre-equilibrated Ni-NTA agarose column and washed with a wash buffer containing 6M GuHCl and 10 mM imidazole at pH 7.5 to remove any non-specifically bound proteins. Subsequently, the protein bound to the Ni-NTA agarose column was eluted with the elution buffer comprising 6M GuHCl and 250 mM imidazole at pH 7.5. The eluted fractions were analyzed by SDS-PAGE for protein homogeneity.

### 2.4. *In vitro* aggregation of full-length TDP-43 and its C-terminal fragments in the presence of Zn^2+^ or Mn^2+^ metal ions or SO_4_^2-^ ions

#### 2.4.1. Full-length TDP-43 aggregation

The full-length TDP-43 protein recombinantly expressed and purified in the presence of 6M GuHCl, following methods described previously [10,32], was first buffer-exchanged to 10M urea dissolved in either PBS, pH 7.5 or 0.1M sodium acetate buffer, pH 5.0. Subsequently, a thioflavin-T (Th-T) assay was performed to analyze the *in vitro* amyloid-like aggregation of full-length TDP-43 in the presence of Zn^2+^ or Mn^2+^ at pH 7.5 or pH 5.0. For this, 50 µM of soluble monomeric full-length TDP-43 was incubated in aggregation buffer containing 3M urea and 500 µM DTT in either PBS, pH 7.5 or 0.1M sodium acetate buffer, pH 5.0. Protein samples were incubated at 37°C either alone or in the stoichiometric ratios of protein to Zn^2+^ = 1:5 or protein to Mn^2+^ = 1:6. In addition, the full-length TDP-43 aggregation was also analyzed in the presence of 120 mM and 200 mM Na_2_SO_4_. 100 mM Zinc chloride and 100 mM Manganese chloride solutions were prepared in respective aggregation buffers either at pH 7.5 or 5.0 as described above in section 2.4. The samples were added with 400 µM Th-T and incubated at 37°C for 14h with agitation. The amyloid-like growth was monitored by recording Th-T fluorescence emission at 488 nm when excited at 442 nm using fluorescence spectrometry [33–35].

#### 2.4.2. TDP-43^2C^ aggregation

The TDP-43^2C^ protein (aa: 193-414) recombinantly expressed and purified in the presence of 6M GuHCl, following methods described previously [10,17], was first buffer-exchanged to 4M urea dissolved either in PBS, pH 7.5 or 0.1M sodium acetate buffer, pH 5.0. Subsequently, the thioflavin-T (Th-T) assay was performed, as described above for the full-length TDP-43 to analyze the *in vitro* amyloid-like aggregation of TDP-43^2C^ in the presence of Zn^2+^ or Mn^2+^ at pH 7.5 or pH 5.0. For this, 400 µM of soluble monomeric TDP-43^2C^ was incubated at 37°C in the aggregation buffer containing 2.5M urea and 500 µM DTT either in PBS, pH 7.5 or in 0.1M sodium acetate buffer, pH 5.0 either alone or added with metal ions in the stoichiometric ratio of protein to Zn^2+^ = 1:5 and protein to Mn^2+^ = 1:6. In addition, TDP-43^2C^ was also analyzed for aggregation in the presence of 120 mM and 200 mM Na_2_SO_4_ to examine the effect of presence of kosmotropic SO_4_^2-^ ions.

#### 2.4.3. TDP-43-LCD aggregation

The TDP-43’s low complexity domain (LCD) (aa: 274-414) recombinantly expressed and purified in presence of 6M GuHCl, following methods similar to as used previously for TDP-43^2C^ [10,28] **(Supplementary figure 1S)**, was dialyzed first against PBS, pH 7.5 for 12h at 4°C and then fresh PBS, pH 7.5 buffer was changed and dialysis was continued further for 12h at 4°C. Upon the completion of dialysis, the TDP-43-LCD was observed to precipitate in the dialysis bag. The precipitated TDP-43-LCD protein was subsequently dissolved using 6M urea in 50 mM phosphate buffer, pH 7.5. Then, thioflavin-T (Th-T) assay was performed, to analyze the *in vitro* amyloid-like aggregation of TDP-43-LCD. For this, 15 µM of soluble monomeric TDP-43-LCD was incubated at 37°C either in the aggregation buffer containing final 1.5M urea alone or added with Zn^2+^ in the stoichiometric ratio of protein to Zn^2+^ = 1:5. The Th-T fluorescence emission intensity was recorded at 488 nm upon excitation at 442 nm at 0h and 6h. To assess the amyloid nature, fold-change in the Th-T emission fluorescence intensity was calculated with respect to the corresponding blank solutions incubated with similar Th-T concentrations. In addition, the aggregation of TDP-43-LCD with or without Zn^2+^ was also assessed by turbidity assay by recording the absorbance at 450 nm at 0h and 6h.

### 2.5 *In vitro* Alexa Flour labelling of TDP-43-LCD and phase separation visualization by fluorescence microscopy in presence of Zn^2+^ ions

For fluorescent labelling, a 50 μL aliquot of TDP-43-LCD protein (215 μM) at PBS pH 7.5 containing 6M GuHCl, was added with 320 μM Alexa Fluor^TM^ 633 NHS ester dye (protein : dye = 1 : 1.5), which covalently attaches to the lysine residues, and the mixture was incubated in dark at room temperature for 2h with gentle intermittent vortexing every 10 min [17,36]. The excess un-reacted dye was removed by dialysis against PBS, pH 7.5 for 36h, with a fresh buffer change every 12h. After the dialysis, the protein was observed to precipitate which was then dissolved in 6M urea in 50 mM phosphate buffer, pH 7.5. For the phase separation visualization, the Alexa Fluor^TM^ 633-labeled and unlabeled TDP-43-LCD proteins were mixed in the ratio of 1 : 15. The aggregation condition, as described above for the turbidity assay (15 μM protein, 1.5M urea, 50 mM phosphate buffer, pH 7.5), was used to examine for the presence of any phase-separated structures of TDP-43-LCD in or absence or presence of Zn^2+^ (protein : Zn^2+^ = 1 : 5) under RFP filter or bright-field in a Leica DM2500 fluorescence microscope. The images were visualized and acquired using 20x objective lens. All the images were background-subtracted and processed for color using ImageJ [37].

### 2.6 Fluorescence microscopy of thioflavin-T-bound aggregates of full-length TDP-43 and its C-terminal fragments

The *in vitro*-made aggregates of full-length TDP-43 and TDP-43^2C^ protein, as described in section 2.4, obtained in the absence or presence of Zn^2+^, Mn^2+^ or SO_4_^2-^ ions were also further examined under the GFP filter using the Leica DM2500 fluorescence microscope and thioflavin-T-bound aggregate images were acquired as per the method described earlier [17]. The fluorescent images were acquired using the 10x objective lens and the obtained images were processed for color and background-subtracted using ImageJ software [37].

### 2.7 Fluorescence spectrometry of deep-blue autofluorescence (dbAF) of aggregates of TDP- 43^2C^

The TDP-43^2C^ was incubated for aggregation in the aggregation buffers as described above in section 2.4 but without the addition of the Th-T dye. The samples were incubated at 37°C for 16h with rotation at 200 rpm either in the presence or absence of Zn^2+^ or Mn^2+^ions. To record deep-blue autofluorescence (dbAF) spectra, the samples were placed in a 10 mm quartz cuvette and excited at 375 nm [24] and emission fluorescence was recorded from 400-600 nm in steps of 10 nm using Spectramax M5e multimode microplate reader. For controls, blank samples lacking the protein but containing everything else, were also simultaneously analyzed for comparison.

### 2.8 Fluorescence microscopy of deep-blue autofluorescence (dbAF) of aggregates of full-length TDP-43 and its C-terminal fragments

As the aggregates of full-length TDP-43 and its C-terminal fragments (TDP-43^2C^ and TDP-43-LCD) exhibited enhanced intensities of deep-blue autofluorescence (dbAF) spectra (400-600 nm) in presence of Zn^2+^ and Mn^2+^ upon excitation at 375 nm, thus, we furthermore examined if the dbAF of the aggregates can be visualized in fluorescence microscopy under the UV filter (excitation filter-360/40 nm, emission filter - 470/40 nm). For this, soluble full-length TDP-43 and its CTFs were incubated in their respective aggregation buffers but without Th-T dye as mentioned above in section 2.6. The samples were incubated either in the absence or presence of Zn^2+^ or Mn^2+^ at 37°C with rotation at 200 rpm. The 5 µL aliquots of samples were taken out at different time points (0h, 6h and 16h) and placed on glass slides and then visualized under the UV filter using the 10x objective lens of Leica DM2500 fluorescence microscope and imaged with an exposure time of 3s for the samples containing full-length TDP-43 and TDP-43^2C^ and 100 ms for samples with TDP-43-LCD. Similarly, the full-length TDP-43 and TDP-43^2C^ samples were also incubated either in the absence or presence of 120 mM and 200 mM Na_2_SO_4_ dissolved in PBS, pH 7.5 and visualized under the UV filter using the 10x objective lens of Leica DM2500 fluorescence microscope. The acquired images were background-subtracted and processed for color using the ImageJ software[37].

### 2.9 Atomic force microscopy

The TDP-43^2C^ protein was incubated for one hour at 37°C with rotation at 200 rpm in the aggregation buffer (2.5M urea, PBS, pH 7.5) either in the absence or presence of Zn^2+^ or Mn^2+^ ions (protein : Zn^2+^ = 1:5 and protein : Mn^2+^ = 1:6). Then, 10 µL aliquots of the samples were placed on freshly cleaved mica sheets and allowed for adsorption for 30 min at room temperature. Subsequently, the mica sheets were washed with water several times and allowed to air-dry at room temperature. The dried mica sheets with adsorbed samples were imaged in tapping mode using an AC160TS cantilever in a multimodal scanning probe microscope (Park Systems) similar to as described previously [32,33]. Imaging was performed with a scan rate of 1 Hz. Analysis of the AFM images was carried out using the Gwyddion software [38]. Aggregate heights vs aggregate count profile was obtained and the normalized height distribution was fit using the “Gaussian fit” function. Furthermore, the mean height of the particles in the images and the standard deviation were obtained using the GraphPad Prism software [39].

## 3. Results and Discussion

### 3.1. Zn^2+^ and Mn^2+^ ions enhance the *in vitro* aggregation of full-length TDP-43 and TDP-43^2C^ at pH 7.5

As the levels of several metals were found significantly elevated in the ALS patients [13] and increased oxidative stress was found also associated with elevated levels of the Zn^2+^, Cu^2+^ and Mn^2+^ metal ions in the spinal cord of TDP-43^A315T^ transgenic mice [14], therefore, we first examined the effect of, yet poorly explored, metal ions on TDP-43’s *in vitro* aggregation. As the full-length TDP-43 protein can undergo proteolytic cleavage into C-terminal fragments (CTFs) in ALS and FTLD patients [4], we first examined the effect of Zn^2+^ and Mn^2+^ ions on the *in vitro* aggregation of recombinantly purified full-length TDP-43 and TDP-43^2C^ (aa: 193-414) that encompasses RRM2 and C-terminal domain (CTD) and bears similarity to a caspase-cleaved fragment that is found in the ALS patients **(Figure 1A)** [4]. The full-length TDP-43 protein sequence has many residues with capability of interaction with Zn^2+^ such as Asp, Glu, Cys and His and some of these residues are also present in the sequence of the CTF, TDP-43^2C^.

**Figure 1.**
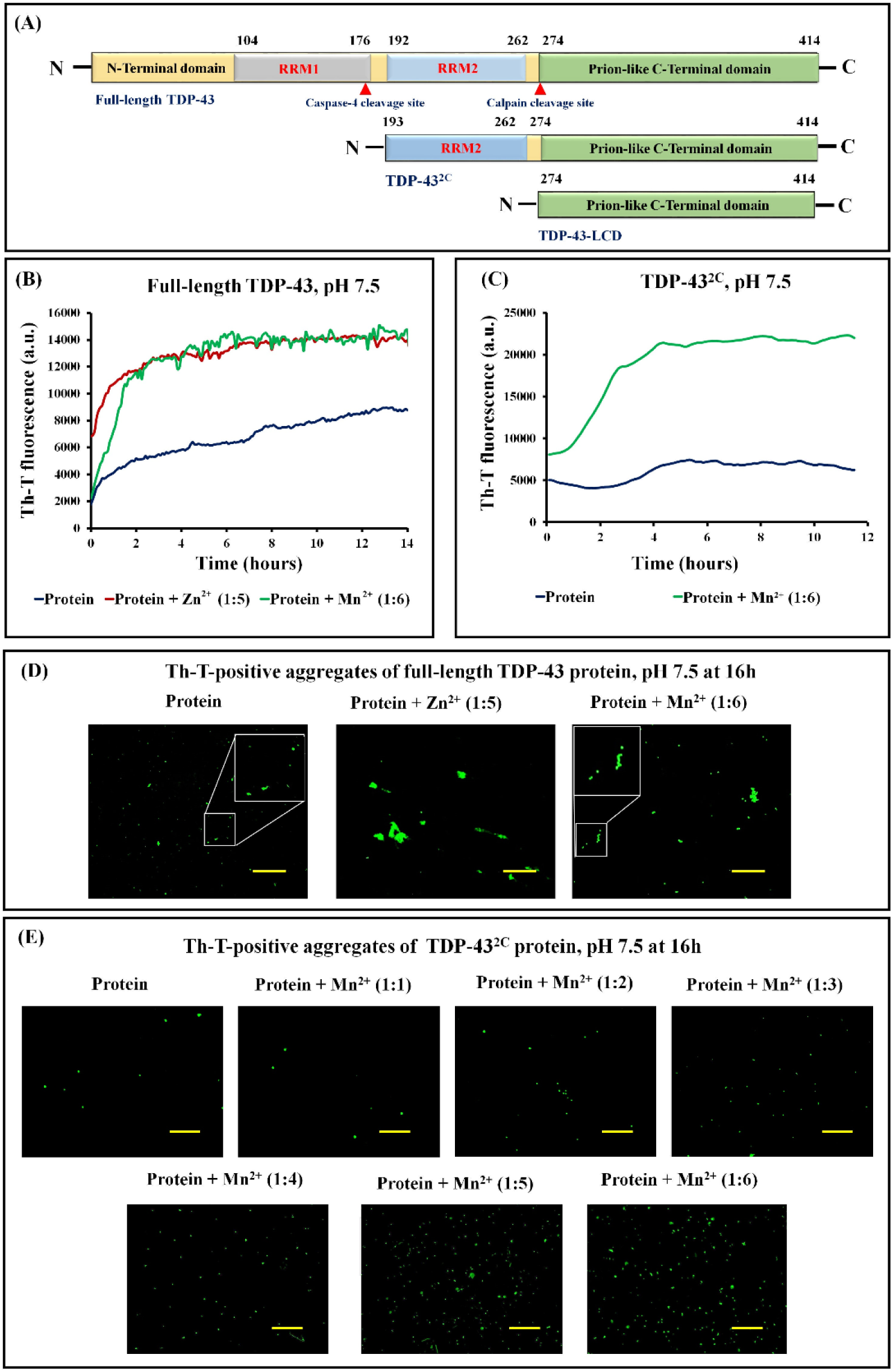
Aggregation and visualization of thioflavin-T-positive aggregates of full-length TDP-43 and TDP-43^2C^ incubated in presence of Zn^2+^ and Mn^2+^ ions at pH 7.5. (A) Schematic representation of the domain organization of the full-length TDP-43 and its C-terminal fragments (CTFs) used in this study. The full-length TDP-43 comprises of an N-terminal domain (aa: 1-89), RRM12 domains (aa: 104-262) and a C-terminal low complexity domain (LCD) (aa: 274-414). A CTF (aa: 193-414) of TDP-43 termed, TDP-43^2C^, encompasses the domains, RRM2 and LCD. Another CTF, TDP-43-LCD, contains only the intrinsically disordered C-terminal low complexity domain (aa: 274-414). (B) Aggregation of full-length TDP-43 in absence or presence of Zn^2+^ and Mn^2+^, monitored by thioflavin-T (Th-T) fluorescence spectroscopy. The full-length TDP-43 protein (50 µM) was incubated in aggregation buffer pH 7.5 either in absence or presence of Zn^2+^ and Mn^2+^ ions at a stoichiometric ratio of 1:5 and 1:6 respectively for 14h and Th-T fluorescence intensity was recorded. The Th-T fluorescence kinetics for the aggregation of the full-length TDP-43 were obtained after subtracting the respective control blanks lacking proteins but containing either only the aggregation buffer or the aggregation buffer added with Zn^2+^ or Mn^2+^ ions. (C) Aggregation of the CTF, TDP-43^2C^, in absence or presence of Mn^2+^ monitored by Th-T fluorescence spectrometry. The TDP-43^2C^ protein (100 µM) was incubated in aggregation buffer pH 7.5 either in absence or in presence of Mn^2+^ ions at a stoichiometric ratio of 1:6 for 12h and Th-T fluorescence was recorded. The Th-T fluorescence kinetics for the aggregation of TDP-43^2C^ were obtained after subtracting the respective control blanks lacking proteins but containing either only the aggregation buffer or the aggregation buffer added with Mn^2+^ ions. (D) Visualization of the Th-T-positive green fluorescent aggregates of full-length TDP-43 obtained in absence or presence of Zn^2+^ and Mn^2+^ after 16h of incubation in aggregation buffer pH 7.5. For recording the fluorescence images, the GFP filter of Leica DM2500 fluorescence microscope and 10x objective lens were used followed by color processing and background subtraction of the images using ImageJ. Scale bar is 200 µm. (E) Visualization of the Th-T-positive green fluorescent aggregates of full-length TDP-43 obtained in absence or presence of different stoichiometric ratios of Mn^2+^ (protein: Mn^2+^ = 1:1-1:6) after 16h of incubation in aggregation buffer pH 7.5. For the recording the fluorescence images, the GFP filter of Leica DM2500 fluorescence microscope and 10x objective lens were used followed by color processing and background subtraction of the images using ImageJ. Scale bar is 200 µm.

Previously, Zn^2+^ was found to enhance the *in vitro* aggregation of recombinantly purified RRM1-2 domain [15] and TDP-43^2C^ CTF [17], however, whether Zn^2+^ can also influence the *in vitro* aggregation of the full-length TDP-43 protein, remains to be examined. Therefore, when we incubated the full-length TDP-43 protein, with a 1 : 5 ratio of protein : Zn^2+^, and examined amyloid-like aggregation by Th-T fluorescence, a four-fold increase in the Th-T fluorescence intensity was observed, after 14h of incubation under agitated conditions, as opposed to only two-fold Th-T fluorescence intensity increase when the full-length TDP-43 was incubated similarly but without the addition of Zn^2+^ (**Figure 1B**). Notably, the Th-T fluorescence intensity increase was also very rapid in the presence of the Zn^2+^ ions (**Figure 1B**). This significant enhancement of the Th-T fluorescence intensity suggests that Zn^2+^ induces and promotes amyloid-like *in vitro* aggregation of the full-length TDP-43 [34].

Similarly, as the full-length TDP-43’s amino acid sequence also has many possible Mn^2+^- interacting residues such as Asp, His and Glu and some of the residues reside in the TDP-43^2C^ region as well, and as so far no studies have examined the effect of Mn^2+^ on the *in vitro* aggregation of full-length TDP-43, therefore we next examined the effect of addition of Mn^2+^ ions on the aggregation of full-length TDP-43 as well as on the TDP-43^2C^ CTF using 1 : 6 (protein : Mn^2+^) stoichiometry. In contrast to the instantaneous Th-T fluorescence intensity increase observed in the presence of Zn^2+^ ions, the incubation of full-length TDP-43 protein with Mn^2+^ resulted only in a steady Th-T fluorescence intensity increase (**Figure 1B**). Strikingly however, a seven-fold increase in the Th-T fluorescence intensity was achieved upon 14h of incubation thereby strongly suggesting an amyloid-like aggregation of the full-length TDP-43 protein in the presence of Mn^2+^ ions [34] (**Figure 1B**).

In the view that the Zn^2+^ and Mn^2+^ ions caused significant enhancement of the Th-T fluorescence emission intensity when incubated with full-length TDP-43, in order to directly visualize the effects and the Th-T-positive aggregates, we also imaged the aggregates under fluorescence microscope using the GFP filter. We find that after 16h of incubation without any added metal ions, the full-length TDP-43 samples manifested dispersed low-intensity green fluorescent aggregates with irregular margins. In comparison, when incubated in the presence of the 1 : 5 ratio of protein to Zn^2+^, the full-length TDP-43 protein samples mostly exhibited relatively larger-sized, irregular-shaped and brighter green fluorescent aggregates suggesting a more efficient Th-T binding and an agglomeration of the aggregates. Notably, in the samples containing the 1 : 6 ratio of full-length TDP-43 to Mn^2+^, we also observed irregular-shaped bright Th-T-positive fluorescent aggregates that were relatively larger than the aggregates of full-length TDP-43 obtained when incubated in absence of metal ions. However, the Mn^2+^-induced full-length TDP-43 aggregates were comparatively smaller than those observed in the presence of Zn^2+^ ions suggesting lesser extent of aggregate agglomeration (**Figure 1D** and **Supplementary figure 2S**). Taken together, the data suggest that while both the Zn^2+^ and Mn^2+^ ions can enhance the amyloid-like aggregation of full-length TDP-43, the resultant aggregates have different levels of agglomeration possibly reflecting mechanistic differences in the effects of these two metal ions.

In view that the Zn^2+^ ions were previously documented to enhance the *in vitro* phase separation and aggregation of the CTF, TDP-43^2C^ [17], we further examined if Mn^2+^ ions can also influence the *in vitro* aggregation of TDP-43^2C^. When TDP-43^2C^ was incubated with an 1 : 6 stoichiometric ratio of protein to Mn^2+^, we observed a sigmoidal aggregation kinetics when monitoring the Th-T fluorescence intensity and an overall ∼ 2.7-fold increase in the Th-T fluorescence intensity over 12h incubation in contrast to ∼1.25-fold increase in case of the TDP-43^2C^ sample incubated similarly but without any metal ions (**Figure 1C**). Furthermore, when the 16h Th-T-positive samples of TDP-43^2C^ protein treated with increasing protein to Mn^2+^ stoichiometric ratios of 1:1 to 1:6, were visualized in the fluorescence microscope using the GFP filter, the sizes and morphologies of the aggregates appeared mostly globular and minute with only a few occasional irregular aggregates similar to as observed in the TDP-43^2C^ alone samples here and as also documented previously [17]. Strikingly however, the increasing levels of Mn^2+^ ions incrementally enhanced the numbers of the Th-T-positive fluorescent aggregates of the TDP-43^2C^ protein (**Figure 1E** and **Supplementary figure 3S**). Overall, while the Mn^2+^ ions can enhance the aggregation of TDP-43^2C^ protein, induction of agglomeration of these aggregates is absent as compared to the Zn^2+^ ion-mediated effects.

### 3.2. Enhanced aggregations of full-length TDP-43 and TDP-43^2C^ due to Zn^2+^ and Mn^2+^ at pH 7.5 also enhance deep-blue autofluorescence

Aggregates of certain proteins including insulin and lysozyme were found to emit deep-blue autofluorescence (dbAF) when excited at certain wavelength in the near-UV region [40]. So far, dbAF generation upon the aggregation of full-length TDP-43 has not been reported. As the presence of Zn^2+^ and Mn^2+^ enhanced the aggregations of full-length TDP-43 and its CTF, TDP-43^2C^ (**Figure 1B** and **1C**), we next examined if the metal ion-induced enhanced aggregations of these proteins also concurrently generate or enhance dbAF. For this, when TDP-43^2C^ samples were incubated for 16h at pH 7.5 for aggregation in the presence of the Zn^2+^ or Mn^2+^ ions and subsequently used to record fluorescence emission (400 – 600 nm) upon excitation at 375 nm, significantly enhanced fluorescence intensities with peak around 440 nm, consistent with dbAF emission, were observed compared to the protein samples incubated in absence of the Zn^2+^ and Mn^2+^ ions (**Figure 2**).

**Figure 2.**
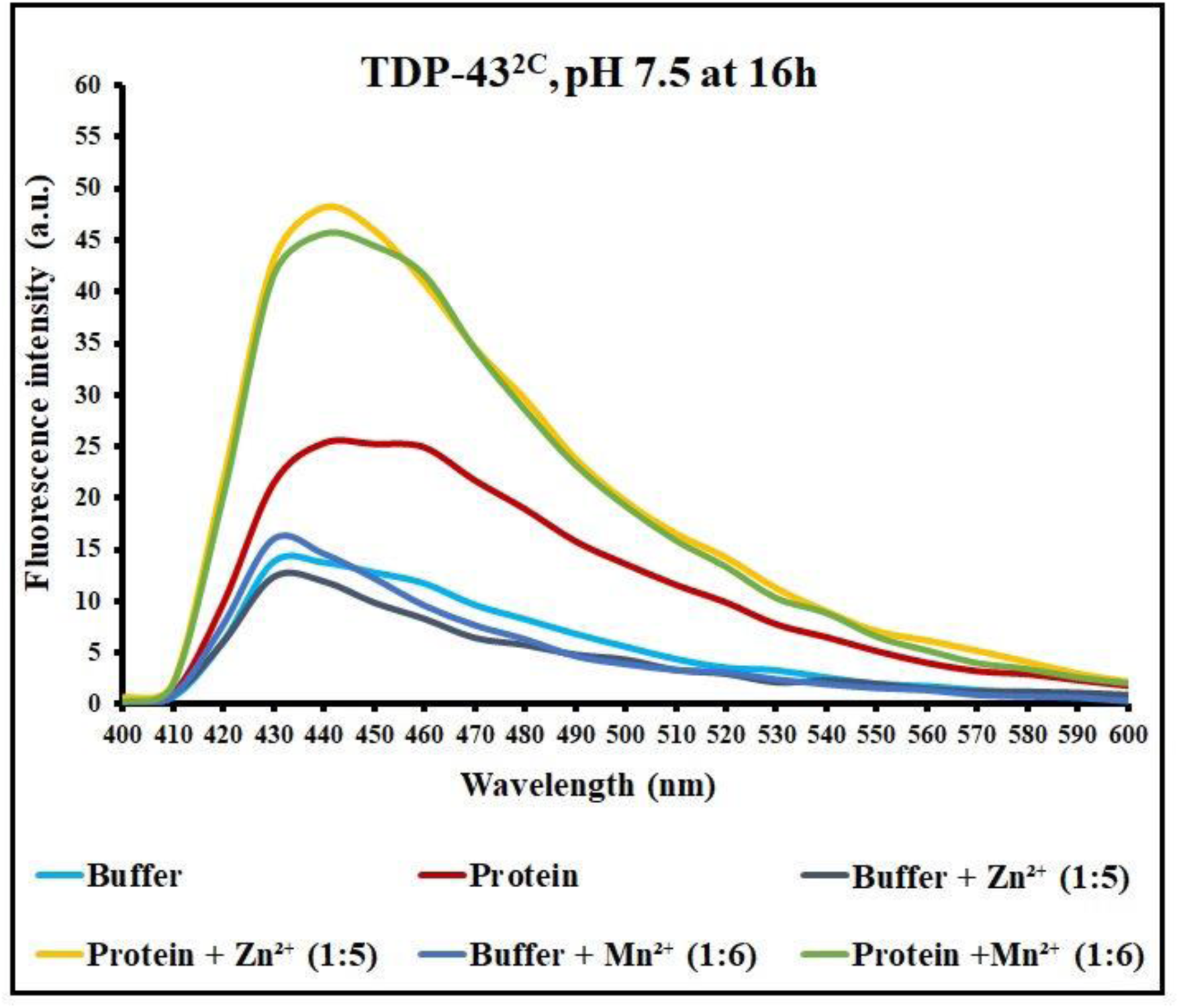
Fluorescence spectrometry of TDP-43^2C^ to detect deep-blue autofluorescence generation upon incubation with Zn^2+^ and Mn^2+^ at pH 7.5. Fluorescence spectra of TDP-43^2C^ aggregated at pH 7.5 in absence or presence of the Zn^2+^ and Mn^2+^ ions to monitor intrinsic dbAF upon excitation at 375 nm. The TDP-43^2C^ protein (400 µM) was incubated for 16h in the aggregation buffer pH 7.5 either in absence or presence of Zn^2+^ and Mn^2+^ at the stoichiometric ratio of 1:5 and 1:6 respectively and the samples were excited at 375 nm and emission fluorescence was recorded from 400-600 nm. The fluorescence spectra were also collected for the respective blanks containing either only the aggregation buffer or the aggregation buffer added with the Zn^2+^ and Mn^2+^ ions.

As the Zn^2+^ and Mn^2+^ ions, caused enhancement of dbAF of TDP-43^2C^ when examined by fluorescence spectrometry, in order to directly visualize the effects and the dbAF-positive aggregates, we also observed the aggregates of both full-length TDP-43 and TDP-43^2C^ under fluorescence microscope using the UV filter. At the zeroth hour of incubation, the full-length TDP-43 protein lacking any added metal ions did not exhibit any aggregates emitting dbAF whereas, the samples treated with Zn^2+^ and Mn^2+^ ions manifested a few minute aggregates with irregular margins emitting dbAF (**Figure 3A** and **Supplementary figure 4S**). When these full-length TDP-43 samples were again visualized after six hours of incubation, the samples incubated without metal ions continued to lack any aggregates emitting dbAF whereas the number of aggregates emanating dbAF were observed to increase in the samples incubated with Zn^2+^ and Mn^2+^ compared to their zero hour incubation (**Figure 3A** and **Supplementary figure 4S**). Strikingly, when examined after 16h incubation, even the full-length TDP-43 samples, incubated without any metal ions, also manifested irregular margin aggregates that emitted dbAF and the numbers of the dbAF-emanating aggregates were furthermore increased in the full-length TDP-43 samples incubated with the Zn^2+^ and Mn^2+^ ions (**Figure 3A** and **Supplementary figure 4S**). Strikingly, unlike the observations on the full-length TDP-43, the TDP-43^2C^ sample incubated in the absence of any added metal ions, did not manifest aggregates emitting dbAF in fluorescence microscopy even after 16 hours of incubation (**Figure 3B** and **Supplementary figure 5S**). However, the TDP-43^2C^ samples incubated with Zn^2+^ or Mn^2+^ ions displayed aggregates with irregular margins that emitted dbAF even at the zeroth hour, and the number of these aggregates were observed to furthermore increase at the sixth and 16^th^ hours of the incubation period (**Figure 3B and Supplementary figure 5S).**

**Figure 3.**
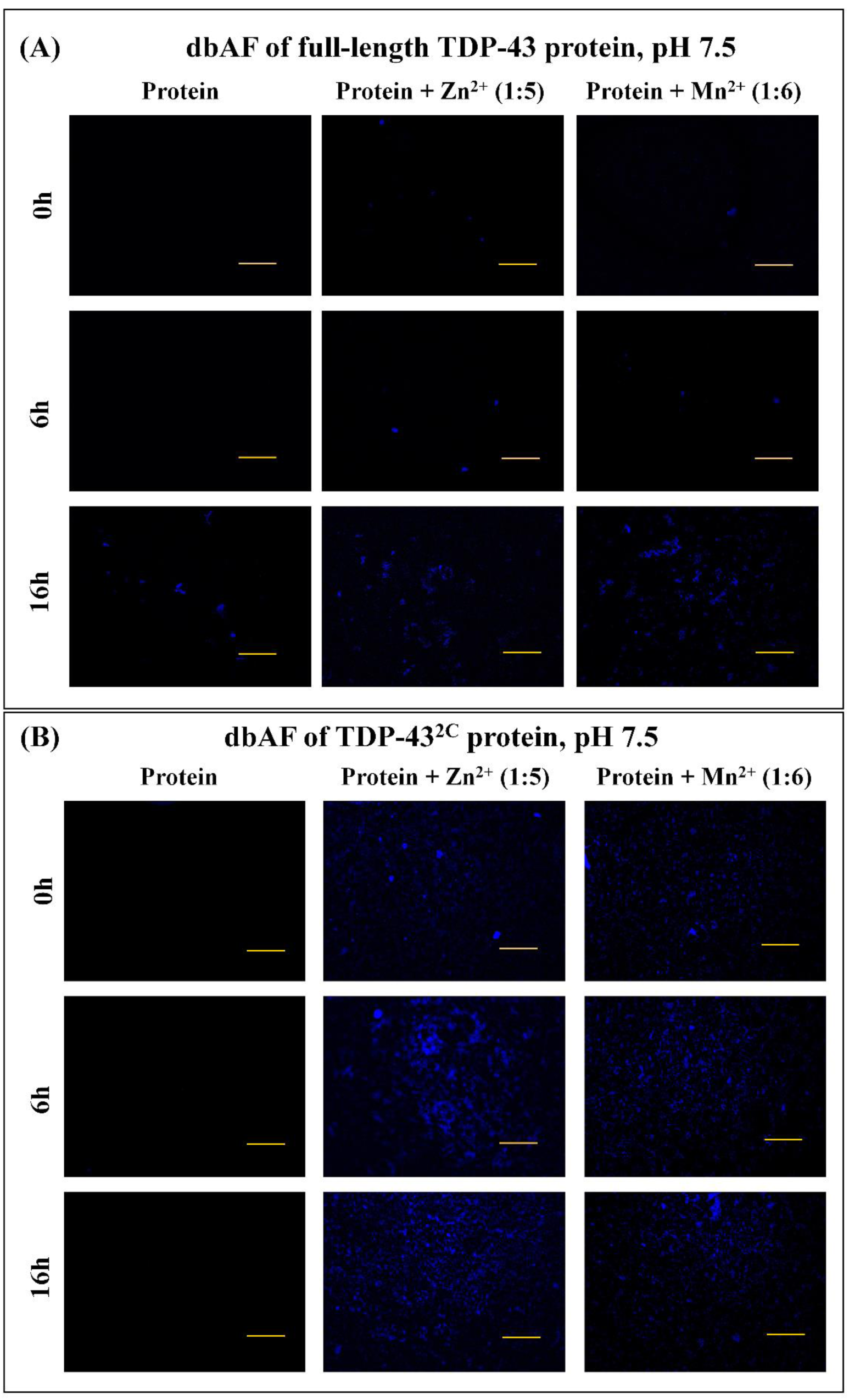
Fluorescence microscopy of full-length TDP-43 and TDP-43^2C^ to detect deep-blue autofluorescence (dbAF) generation upon incubation with Zn^2+^ and Mn^2+^ at pH 7.5. **(A)** Fluorescence micrographs of deep blue autofluorescence (dbAF)-positive fluorescent aggregates of full-length TDP-43 obtained in absence or presence of Zn^2+^ and Mn^2+^ at 0h, 6h and 16h of incubation in the aggregation buffer pH 7.5. For recording the fluorescence images, the UV filter of Leica DM2500 fluorescence microscope, 10x objective lens and imaging exposure time of 3s, were used followed by color processing of the images using ImageJ. Scale bar is 200 µm. **(B)** Fluorescence micrographs of dbAF-positive fluorescent aggregates of TDP-43^2C^ obtained in absence or presence of Zn^2+^ and Mn^2+^ at 0h, 6h and 16h of incubation in aggregation buffer pH 7.5 For recording the fluorescence images, the UV filter of Leica DM2500 fluorescence microscope, 10x objective lens and imaging exposure time of 3s, were used followed by color processing of the images using ImageJ. Scale bar is 200 µm.

Consistent with the acceleration of the aggregation of TDP-43^2C^ and the concurrent dbAF emission being due to the aggregation-enhancing effects of the metal ions, when in a control experiment, Zn^2+^ ion-treated TDP-43^2C^ sample at pH 7.5 was also added with molar excess of the metal chelator molecule, EDTA, (TDP-43^2C^ : Zn^2+^ : EDTA = 1 : 5 : 10) at the zero hour and the samples were examined after 16h incubation, as expected, any TDP-43^2C^ aggregates manifesting dbAF were not observed (**Supplementary figure 6S)**.

Taken together, the fluorescence spectrometry and fluorescence microscopy indicate that the aggregates of full-length TDP-43 and TDP-43^2C^ can emit dbAF and the acceleration and enhancement of the aggregation of these proteins in the presence of the Zn^2+^and Mn^2+^ ions can concomitantly also enhance the emission of dbAF, which could potentially be used as a tool to detect the TDP-43 aggregation in a label-free fashion.

### 3.3. Zn^2+^, but not Mn^2+^, can enhance the *in vitro* aggregations of full-length TDP-43 and TDP-43^2C^ at pH 5.0

As the abilities of some of the amino acids to interact with Zn^2+^ or Mn^2+^ ions can be affected by pH conditions due to their side chain pKa [41–43], we next examined if the effect of these metal ions on the *in vitro* aggregation of full-length TDP-43 and TDP-43^2C^, when examined at an acidic pH 5.0, is alike to or different from the observations at pH 7.5.

Similar to as described in section 3.1 (**Figure 1B**), when we monitored the aggregation kinetics of full-length TDP-43 for 14 hours by monitoring Th-T fluorescence intensity by incubating the protein at pH 5.0 in the absence or presence of molar excess of Zn^2+^ and Mn^2+^ ions, we observed that Zn^2+^, but not the Mn^2+^ ions, caused significant acceleration and enhancement of the Th-T fluorescence intensity thereby suggesting increased aggregation of full-length TDP-43 (**Figure 4A**). Furthermore, a direct visualization of the Th-T fluorescence-positive assemblies using the GFP filter in fluorescence microscopy also revealed the presence of relatively more number and larger-sized aggregates of full-length TDP-43 in the Zn^2+^-treated samples as compared with the untreated or Mn^2+^-treated samples thereby confirming the Zn^2+^-mediated enhancement of the aggregation even at pH 5.0 (**Figure 4B** and **Supplementary figure 7S**).

**Figure 4.**
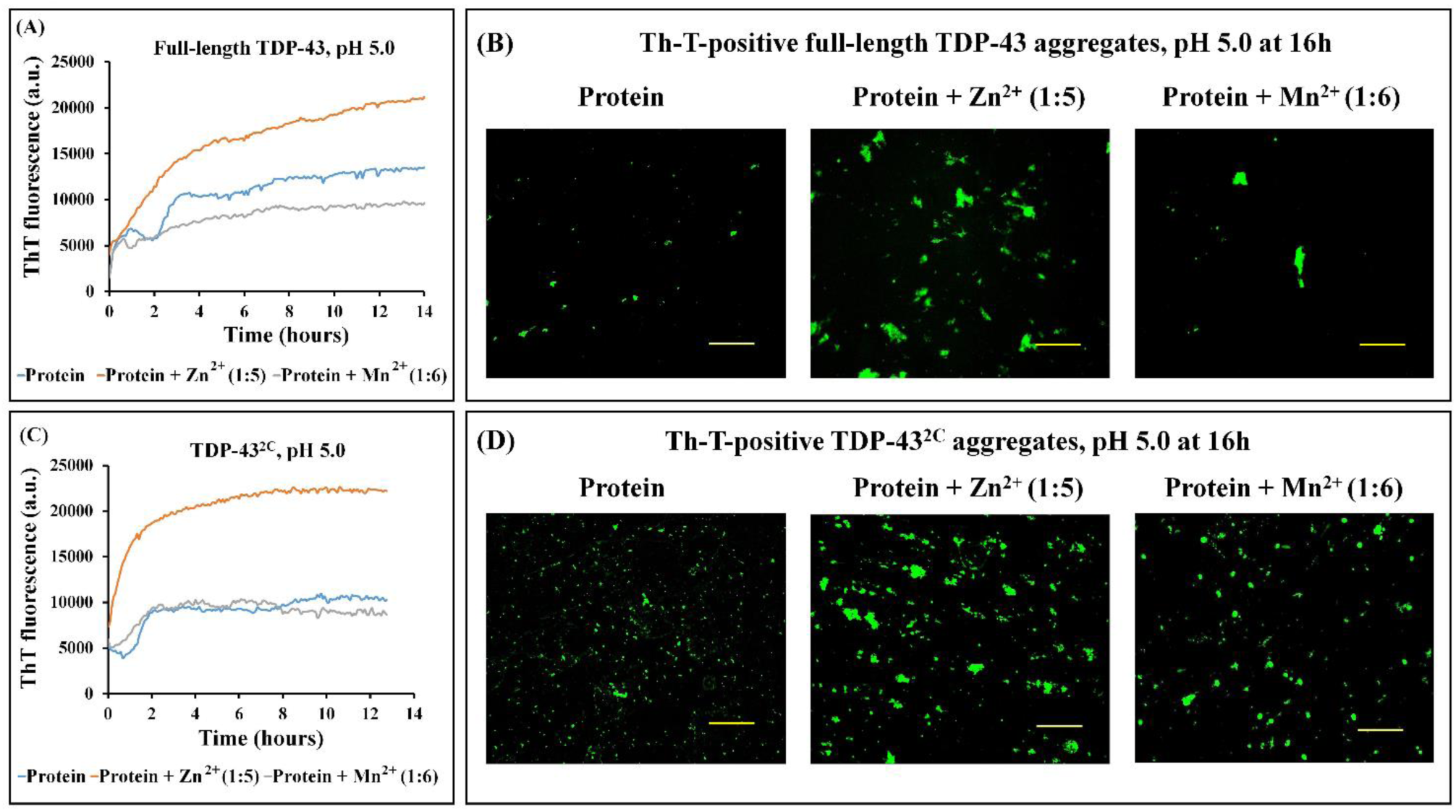
Aggregation and visualization of thioflavin-T-positive aggregates of full-length TDP-43 and TDP-43^2C^ upon incubation in presence of Zn^2+^ and Mn^2+^ at pH 5.0. **(A)** Aggregation of full-length TDP-43 in absence or presence of the Zn^2+^ and Mn^2+^ ions monitored by thioflavin-T (Th-T) fluorescence spectrometry. The full-length TDP-43 protein (50 µM) was incubated in aggregation buffer pH 5.0 either in absence or in presence of the Zn^2+^ and Mn^2+^ ions at a stoichiometric ratio of 1:5 and 1:6 respectively for 14h and Th-T fluorescence was recorded. The Th-T fluorescence kinetics for the aggregation of the full-length TDP-43 were obtained after subtracting the respective control blanks lacking proteins but containing either only the aggregation buffer or the aggregation buffer added with the Zn^2+^ or Mn^2+^ ions. **(B)** Visualization of the Th-T-positive green fluorescent aggregates of full-length TDP-43 obtained in absence or presence of Zn^2+^ and Mn^2+^ after 16h of incubation in aggregation buffer pH 5.0. For recording the fluorescence images, the GFP filter of Leica DM2500 fluorescence microscope and 10x objective lens were used followed by color processing and background subtraction of the images using ImageJ. Scale bar is 200 µm. **(C)** Aggregation of TDP-43^2C^ in absence or presence of Zn^2+^ and Mn^2+^ monitored by Th-T fluorescence spectrometry. The TDP-43^2C^ protein (400µM) was incubated in aggregation buffer pH 5.0 either in absence or in presence of the Zn^2+^ and Mn^2+^ ions at a stoichiometric ratio of 1:5 and 1:6 respectively for 14h and Th-T fluorescence was recorded. The Th-T fluorescence kinetics for the aggregation of full-length TDP-43 were obtained after subtracting the respective control blanks lacking proteins but containing either only the aggregation buffer or the aggregation buffer added with the Zn^2+^ or Mn^2+^ ions. **(D)** Visualization of the Th-T-positive green fluorescent aggregates of TDP-43^2C^ obtained in absence or presence of Zn^2+^ and Mn^2+^ after 16h of incubation in the aggregation buffer pH 5.0. For recording the fluorescence images, the GFP filter of Leica DM2500 fluorescence microscope and 10x objective lens were used followed by color processing and background subtraction of the images using ImageJ. Scale bar is 200 µm.

Likewise, when we monitored the aggregation kinetics of TDP-43^2C^ for 14 hours by recording Th-T fluorescence intensity upon incubating the protein at pH 5.0 in the absence or presence of molar excess of Zn^2+^ and Mn^2+^ ions, once again we observed that Zn^2+^, but not the Mn^2+^ ions, caused significant acceleration and enhancement of the Th-T fluorescence intensity thereby suggesting increased aggregation of TDP-43^2C^ (**Figure 4C**) similar to as observed for the full-length TDP-43 at pH 5.0. When we visualized the Th-T-positive TDP-43^2C^ aggregates under the GFP filter in fluorescence microscopy, the TDP-43^2C^ samples, incubated similarly in absence of any metal ions at pH 5.0, manifested aggregates with minute sizes (**Figure 4D** and **Supplementary figure 7S**). Strikingly however, the Zn^2+^-treated TDP-43^2C^ samples at pH 5.0 showed a higher number of relatively larger-sized, Th-T-fluorescence positive aggregates with irregular margins confirming the enhancement of the TDP-43^2C^ aggregation even at pH 5.0 similar to as observed previously at pH 7.5 [17] (**Figure 4D** and **Supplementary figure 7S**). Notably, the Mn^2+^-treated TDP-43^2C^ samples, when visualized similarly by fluorescence microscopy, manifested irregularly-shaped Th-T fluorescence-positive aggregates despite that no enhancement in the Th-T fluorescence intensity was observed (**Figure 4C**), which possibly suggests that Mn^2+^ induces subtle changes in the properties of the TDP-43^2C^ aggregates, such as agglomeration or conformational changes, without affecting the overall aggregation levels (**Figure 4D** and **Supplementary figure 7S**).

Taken together, the data suggests that the effects of Mn^2+^ on the aggregations of full-length TDP-43 and TDP-43^2C^ are pH-dependent whereas Zn^2+^ promotes and enhances the aggregations of these proteins at pH 5.0 as well as pH 7.5.

### 3.4. Enhanced aggregations of full-length TDP-43 and TDP-43^2C^ due to Zn^2+^ at pH 5.0 also enhance deep-blue autofluorescence (dbAF) whereas the presence of Mn^2+^ does not

As the Zn^2+^ ions enhanced the Th-T fluorescence-positive aggregations of full-length TDP-43 and TDP-43^2C^ even at pH 5.0, whereas Mn^2+^ failed to enhance the aggregations at pH 5.0, we next examined if these observed effects at pH 5.0 are also consistently relayed towards affecting the generation of the deep-blue autofluorescence (dbAF) by these proteins. For this, when the full-length TDP-43 samples were directly visualized under fluorescence microscope using the UV filter, of the presence of dbAF for the full-length TDP-43 samples pre-incubated in the absence or presence of Zn^2+^ and Mn^2+^ ions at pH 5.0. At the zeroth hour of incubation, the full-length TDP-43 protein sample lacking any metal ions as well as those added with the Mn^2+^ ions, did not exhibit any dbAF-positive aggregates whereas, the Zn^2+^-treated full-length TDP-43 sample manifested a few dbAF-positive aggregates with irregular margins (**Figure 5A** and **Supplementary figure 8S**). After six hours of incubation, the full-length TDP-43 protein sample lacking any metal ions as well as those incubated with the Mn^2+^ ions manifested the appearance of a few irregular margin aggregates with dbAF, whereas the Zn^2+^-treated full-length TDP-43 sample displayed dramatically high number of irregular-shaped aggregates with dbAF (**Figure 5A** and **Supplementary figure 8S**). Upon further examination at 16 hours post-incubation, while the sizes of the aggregates emitting dbAF were observed to increase in the samples lacking any metal ions as well as those containing the Mn^2+^ ions, the overall number of the aggregates were quite significantly lower than the samples incubated with Zn^2+^ (**Figure 5A** and **Supplementary figure 8S**). Overall, while the incubation of full-length TDP-43 at pH 5.0 with Zn^2+^ is observed to dramatically enhance the appearance of aggregates emitting dbAF, in contrast, the presence of Mn^2+^ ions have no such observable effect.

**Figure 5.**
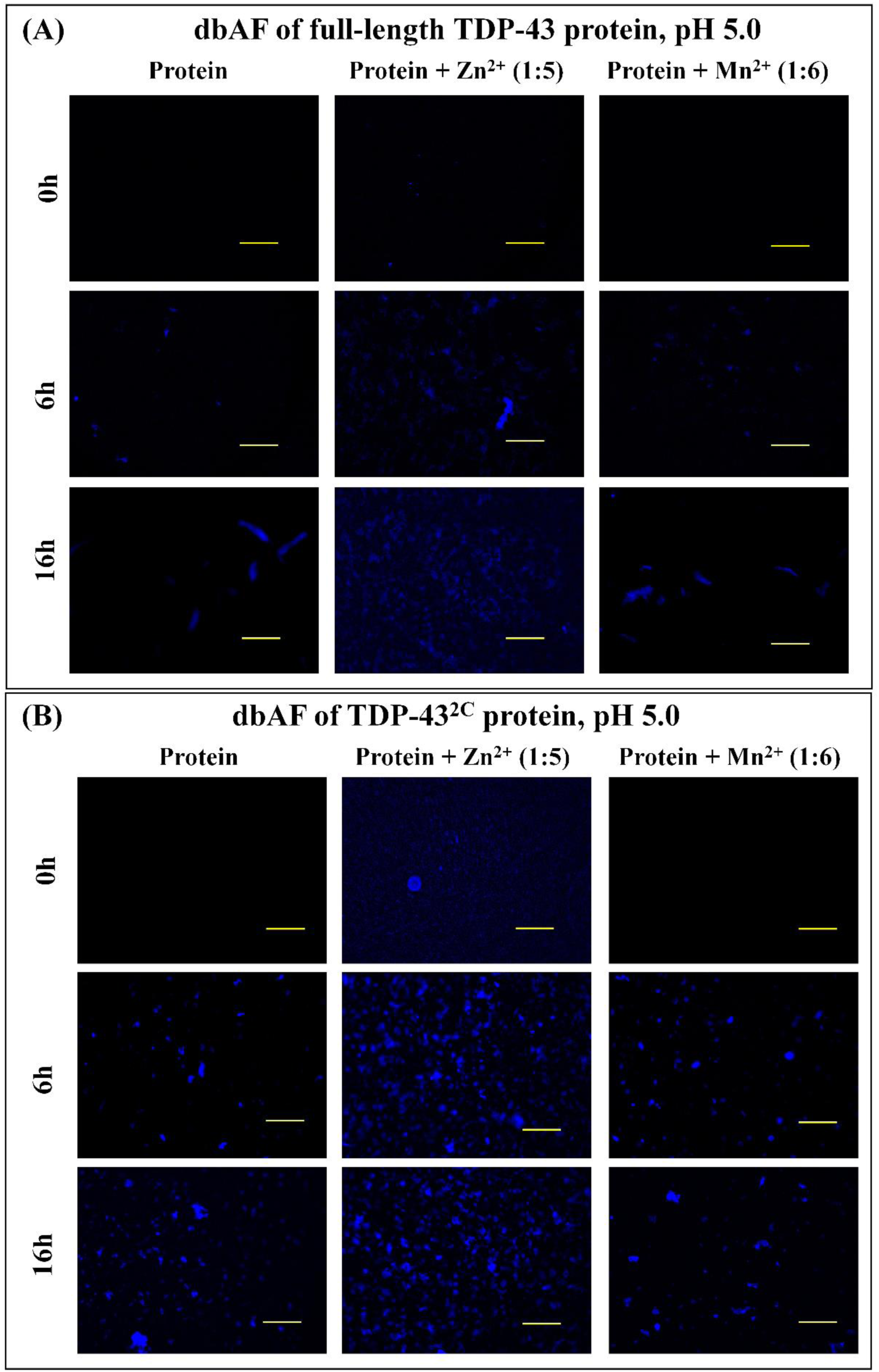
Fluorescence microscopy of full-length TDP-43 and TDP-43^2C^ to detect deep-blue autofluorescence (dbAF) generation upon incubation with Zn^2+^ and Mn^2+^ at pH 5.0. **(A)** Fluorescence micrographs of deep blue autofluorescence (dbAF)-positive fluorescent aggregates of full-length TDP-43 obtained in absence or presence of Zn^2+^ and Mn^2+^ at 0h, 6h and 16h of incubation in aggregation buffer pH 5.0. For acquiring the fluorescence images, the UV filter of Leica DM2500 fluorescence microscope, 10x objective lens and imaging exposure time of 3s, were used followed by color processing of the images using ImageJ. Scale bar is 200 µm. **(B)** Fluorescence micrographs of dbAF-positive fluorescent aggregates of TDP-43^2C^ obtained in absence or presence of Zn^2+^ and Mn^2+^ at 0h, 6h and 16h of incubation in the aggregation buffer pH 5.0. For recording the fluorescence images, the UV filter of Leica DM2500 fluorescence microscope, 10x objective lens and imaging exposure time of 3s, were used followed by color processing of the images using ImageJ. Scale bar is 200 µm.

Next, we also examined the effect of incubation of the TDP-43^2C^ protein with Zn^2+^ and Mn^2+^ at pH 5.0 on the generation of dbAF using the UV filter in fluorescence microscopy. Coherent with the observed lack of enhancement of the aggregation of TDP-43^2C^ at pH 5.0 in the presence of the Mn^2+^ ions (**Figure 4C**), there was no enhancing effect of Mn^2+^ on the dbAF generation by TDP-43^2C^ as compared to the TDP-43^2C^ samples incubated similarly but lacking Mn^2+^. Similar to the untreated samples, the treatment of TDP-43^2C^ at pH 5.0 with Mn^2+^ did not cause appearance of any dbAF-positive aggregates at the zeroth hour of incubation and while some dbAF-positive aggregates appeared at 6h and 16h post-incubation their numbers and the morphological patterns were similar to the untreated TDP-43^2C^ samples thereby confirming the lack of enhancing effect of Mn^2+^ on the TDP-43^2C^ aggregation and dbAF generation at pH 5.0 (**Figure 5B** and **Supplementary figure 9S**). In contrast, upon incubation of TDP-43^2C^ at pH 5.0 with the Zn^2+^ ions, appearance of dbAF-positive aggregates was observed even at zeroth hour of incubation, which furthermore increased in numbers when examined at six and 16 h post-incubation (**Figure 5B** and **Supplementary figure 9S**).

### 3.5. AFM imaging supports enhancement of TDP-43^2C^ aggregation in presence of Zn^2+^ and Mn^2+^ at pH 7.5

As, even at the zeroth hour, we observed an enhancement of the Th-T fluorescence-positive aggregation as well as deep-blue autofluorescence (dbAF) of TDP-43^2C^ in presence of Zn^2+^ and Mn^2+^ at pH 7.5 **(Figure 3B**), therefore, we visualized the TDP-43^2C^ aggregates, incubated for one hour with or without Zn^2+^ and Mn^2+^ at pH 7.5, using atomic force microscope (AFM) imaging. When we analyzed ∼ 426 particles from the AFM images of the TDP-43^2C^ samples incubated without any metal ions, dispersed spherical-shaped particles with an average height of 4.3 ± 0.6 nm were observed **(Figure 6A** and **Supplementary figure 10S)**. Strikingly, in the TDP-43^2C^ samples incubated in presence of Zn^2+^, we observed more and larger particles manifesting an average height of 160.4 ± 39.9 nm when counted among ∼796 particles **(Figure 6B** and **Supplementary figure 10S)**. Also, in presence of Mn^2+^, we observed moderate-sized spherical-shaped particles with an average height of 20.5 ± 6 nm upon analyzing ∼759 particles **(Figure 6C** and **Supplementary figure 10S)**. Overall, the AFM data corroborates that Zn^2+^ and Mn^2+^ ions accelerate and enhance the aggregation of TDP-43^2C^ at pH 7.5 and the presence of aggregates at an early stage is consistent with the manifestation of the early dbAF-positive aggregates in the Zn^2+^ and Mn^2+^-treated TDP-43^2C^ samples at pH 7.5.

**Figure 6.**
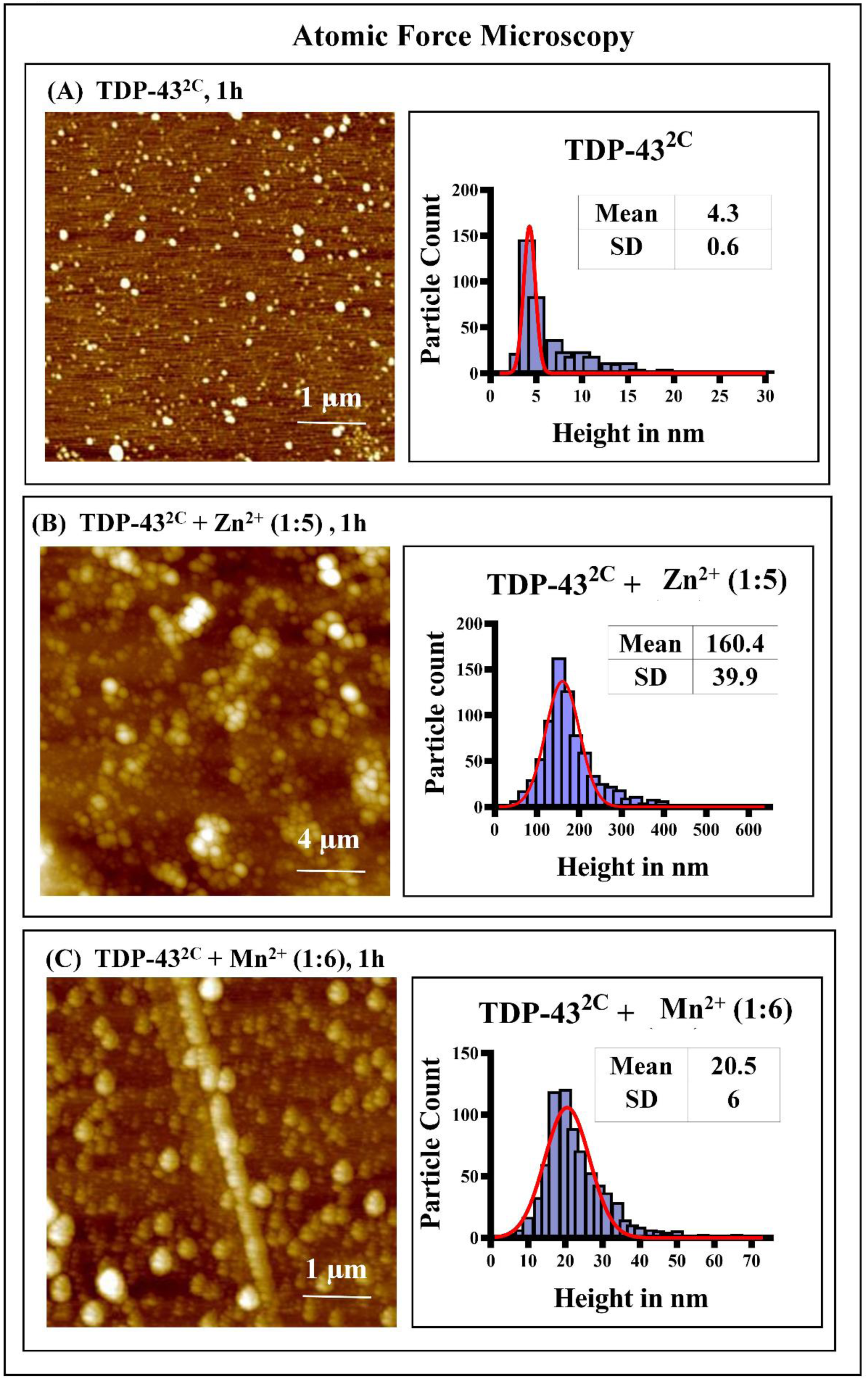
AFM imaging of TDP-43^2C^ incubated in presence of Zn^2+^ and Mn^2+^ at pH 7.5. The morphological characteristics of TDP-43^2C^ aggregated at pH 7.5 either in absence or presence of Zn^2+^ and Mn^2+^ ions were analyzed by AFM imaging. The TDP-43^2C^ protein (400 µM), incubated for one hour at 37°C either in absence or presence of Zn^2+^ and Mn^2+^ ions at respective stoichiometric ratio of 1:5 and 1:6, was drop-casted on freshly cleaved mica sheets then allowed for 30 min adsorption followed by several washes with water and air-drying at room temperature. AFM images were acquired in tapping mode in a multimodal scanning probe microscope (Park Systems) using an AC160TS cantilever. The AFM images were processed and the number of particles were counted using the Gwyddion software. The red curve represents the fit using the Gaussian function in the graph pad prism software. **(A)** AFM image of TDP-43^2C^ incubated for 1h in absence of any added metal ions. Scale bar – 1µm. **(B)** AFM image of TDP-43^2C^ incubated for 1h with added Zn^2+^ ions. Scale bar – 4µm. **(C)** AFM image of TDP-43^2C^ incubated for 1h with added Mn^2+^ ions. Scale bar – 1µm.

### 3.6. Zn^2+^-induced initial phase-separated droplets of TDP-43-LCD do not emit dbAF whereas mature solid-like aggregates of both amyloid and non-amyloid nature manifest dbAF

Among the several C-terminal fragments (CTFs) that can be generated due to proteolytic cleavage of full-length TDP-43, one CTF comprises of only the low-complexity C-terminal domain of TDP-43 (TDP-43-LCD) **(Figure 1A)** [4]. Earlier studies have shown that under different *in vitro* incaution conditions, TDP-43-LCD can undergo liquid-like phase separation into droplets or convert into β-sheet-rich irreversible amyloid-like fibrillar aggregates [7,44].

So far, any effect of Zn^2+^ ions on the *in vitro* phase separation or aggregation of TDP-43-LCD, has not been analyzed. Thus, we examined the *in vitro* aggregation, phase separation and the generation of dbAF by TDP-43-LCD at pH 7.5 in the absence or presence of Zn^2+^ at a stoichiometric ratio of 1 : 5 (protein : Zn^2+^) using turbidity measurements, Th-T fluorescence spectrometry of unlabeled-TDP-43-LCD and fluorescence microscopy of Alexa Fluor-633-labeled-TDP-43-LCD protein.

When TDP-43-LCD protein samples incubated with or without Zn^2+^ were analyzed for formation of amyloid-like Th-T-positive aggregates, the sample lacking Zn^2+^ manifested a striking 15-fold increase in the Th-T fluorescence intensity after 6h incubation thereby strongly indicating the formation of amyloid-like aggregates (**Figure 7A**). In comparison, the TDP-43-LCD sample incubated in the presence of Zn^2+^ showed less than a 5-fold increase in the Th-T fluorescence intensity even after 6h post-incubation thereby indicating a retarding effect of Zn^2+^ on the *in vitro* amyloid-like aggregation of TDP-43-LCD **(Figure 7A)**.

**Figure 7.**
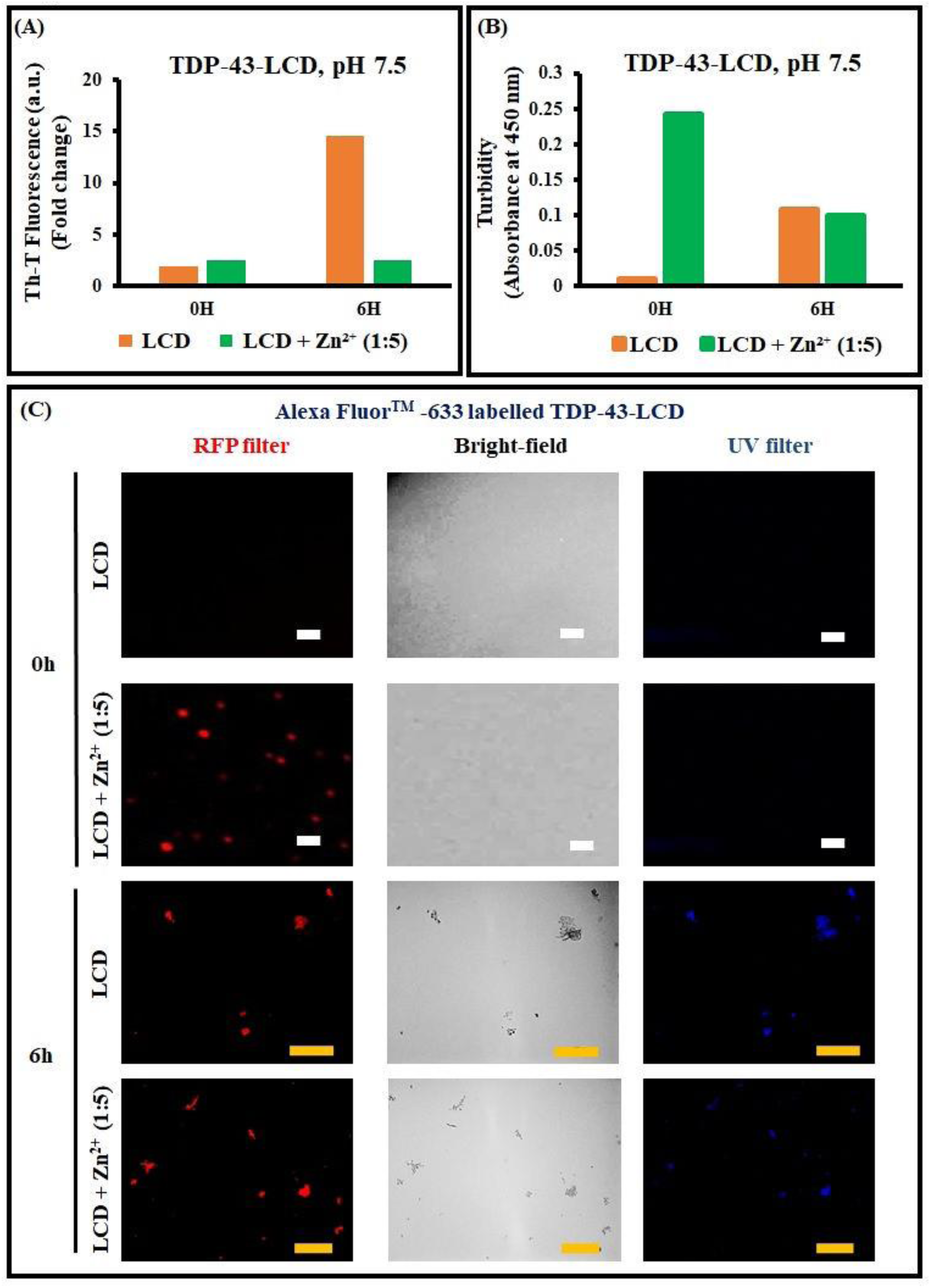
Effect of the Zn^2+^ ions on the *in vitro* aggregation, phase separation and deep-blue autofluorescence (dbAF) of TDP-43-LCD. **(A)** Fold change in the Th-T fluorescence intensity of the TDP-43-LCD samples over time when incubated in absence or presence of the Zn^2+^ ions at pH 7.5. TDP-43-LCD (15 µM) was incubated in 50 mM phosphate buffer, pH 7.5 containing 1.5M urea in absence or presence of added Zn^2+^ ions (protein : Zn^2+^ = 1 : 5) and Th-T fluorescence emission intensity was recorded at 488 nm upon excitation at 442 nm to examine the formation of amyloid-like aggregates. Fold-change in the Th-T fluorescence emission intensity at 488 nm of the samples incubated for 0 and 6h were obtained by comparison with Th-T fluorescence emission intensities of the respective control blank samples lacking the protein and then plotted as a bar chart. **(B)** Solution turbidity of the TDP-43-LCD samples over time when incubated in absence or presence of the Zn^2+^ ions at pH 7.5. The TDP-43-LCD protein samples incubated for 0 or 6h in absence or presence of Zn^2+^ (protein : Zn^2+^ = 1 : 5) were examined for turbidity by recording absorbance at 450 nm. The absorbance recorded at 450 nm were first blank-subtracted and then represented as a bar chart. **(C)** Phase separation and deep-blue autofluorescence (dbAF) generation by TDP-43-LCD labelled with Alexa fluor-633 fluorescent dye. To examine the type of species formed, phase-separated globules or irregular aggregates, TDP-43-LCD labelled with Alexa fluor-633 and unlabelled TDP-43-LCD were mixed in the ratio of 1 : 15 in the aggregation buffer with or without the addition of the Zn^2+^ ions and examined at 0h and 6h of incubation using the RFP filter and 20x objective lens in fluorescence microscopy. To examine the emission of dbAF by the phase-separated globules or the irregular aggregates, the samples examined under the RFP filter were also visualized using the UV filter and 20x objective in the fluorescence microscopy. Acquired images were processed for color using ImageJ. White color scale bars – 10 µm; Yellow color scale bars –100 µm.

Next, we analyzed the effect of Zn^2+^ on TDP-43-LCD *in vitro* aggregation by monitoring the solution turbidity by recording the absorbance at 450 nm at the initial incubation time and 6h post-incubation. Strikingly, TDP-43-LCD incubated in the presence of Zn^2+^ showed an instantaneous burst of high turbidity at the zeroth hour itself which was surprisingly reduced upon furthermore incubation of 6h (**Figure 7B**). In contrast, the TDP-43-LCD sample incubated similarly but lacking the Zn^2+^ ions, did not manifest initial high turbidity although, the turbidity increased upon further 6h incubation and its turbidity reached similar levels to that of the TDP-43-LCD incubated in the presence of Zn^2+^ (**Figure 7B**). This suggests different mechanisms of the TDP-43-LCD aggregation in the presence and absence of the Zn^2+^ ions.

Notably, both the liquid-like phase separation as well as the solid-like aggregation of the TDP-43-LCD protein have been previously documented to enhance the solution turbidity [36,45,46]. As we observed an instantaneous enhancement of the solution turbidity of TDP-43-LCD in presence of Zn^2+^ ions (**Figure 7B**), which was not associated with any concomitant Th-T fluorescence increase (**Figure 7A**), we hypothesized the Zn^2+^ ions-mediated enhancement of turbidity to be a result of potential droplet-like phase separation of TDP-43-LCD at the zeroth hour and the relative reduction in the turbidity at the sixth hour to be possibly a consequence of the conversion and maturation to solid-like TDP-43-LCD aggregates. To test this, when we visualized Alexa fluor-633-labeled-TDP-43-LCD, incubated in absence or presence of Zn^2+^ ions, by fluorescence microscopy under RFP filter, while the samples lacking Zn^2+^ ions did not manifest any fluorescent assemblies at the zeroth hour of incubation, those incubated with the Zn^2+^ ions indeed manifested droplet-like globular-shaped species with smooth margins of ∼ 4 μm diameter thereby suggesting a potentially liquid-like phase separation of Alexa fluor-633-labeled-TDP-43-LCD in presence of Zn^2+^ ions (**Figure 7C** and **Supplementary figure 11S**). Notably, the samples of Alexa fluor-633-labeled-TDP-43-LCD in presence of Zn^2+^ when further incubated for six hours, mostly displayed irregular-margin species of relatively larger sizes thereby confirming the maturation and conversion of the initially observed globular droplets into solid-like aggregates (**Figure 7C** and **Supplementary figure 12S**). Similarly, the Alexa fluor-633-labeled-TDP-43-LCD samples lacking Zn^2+^ ions also manifested irregular-margin species after six hours of incubation suggesting spontaneous solid-like aggregation of Alexa fluor-633-labeled-TDP-43-LCD even in the absence of Zn^2+^ ions (**Figure 7C** and **Supplementary figure 12S**).

The results from the Th-T fluorescence intensity monitoring, turbidimetry and fluorescence microscopy of the Alexa fluor-633-labeled-TDP-43-LCD, all taken together, support that the presence of Zn^2+^ ions cause phase separation of TDP-43-LCD into droplets and while the droplets mature into solid-like aggregates, as observed at six hours post-incubation, they differ from the spontaneously formed aggregates and manifest non-amyloid-like properties. Thus, to furthermore elucidate which of the three types of the TDP-43-LCD species, i.e. phase-separated droplets, solid-like non-amyloid aggregates or amyloid-like aggregates, manifest the deep-blue autofluorescence (dbAF), we also examined the Alexa fluor-633-labeled-TDP-43-LCD protein incubated in absence or presence of the Zn^2+^ ions at 0h and 6h post-incubation, under UV filter in fluorescence microscopy. Notably, the phase-separated droplets of Alexa fluor-633-labeled-TDP-43-LCD did not manifest any dbAF generation but, both the solid-like non-amyloid-like as well as the amyloid-like aggregates manifested dbAF **(Figure 7C** and **Supplementary figures 11S** and **12S)**. As the phase-separated liquid-like droplets of intrinsically disordered proteins are mostly non-pathogenic and even participate in diverse physiological roles [47,48], recording the dbAF could potentially be useful in distinguishing pathogenic protein aggregates from non-pathogenic phase-separated droplets.

### 3.7. Enhancement of full-length TDP-43 and TDP-43^2C^ aggregation induced by the kosmotropic SO_4_^2-^ ions also enhances deep-blue autofluorescence

As metal ion-induced enhancement of the aggregation of full-length TDP-43 and its CTF, TDP-43^2C^ enhanced the deep-blue autofluorescence (dbAF) generation, thus, we checked whether the dbAF generation is a metal-ion dependent phenomenon or any other enhancers of the aggregation can also enhance the dbAF emission from full-length TDP-43 and TDP-43^2C^. Previously, a kosmotropic anion, SO_4_^2-^, was documented to increase the *in vitro* aggregation of the TDP-43^2C^ protein [28] thus, we first examined if SO_4_^2-^ ions can also enhance the aggregation of the full-length TDP-43 and then investigated if the SO_4_^2-^ ion-induced enhanced aggregations of full-length TDP-43 and TDP-43^2C^ are also accompanied by enhanced emission of dbAF.

Consistent with previous reports, the presence of 120 or 200 mM SO_4_^2-^ ions, dramatically increased the sizes of the Th-T fluorescence-positive the TDP-43^2C^ protein aggregates, as observed under the GFP filter of fluorescence microscopy, which were of irregular shapes suggesting solid-like aggregation in presence of the SO_4_^2-^ ions in contrast to the non-treated TDP-43^2C^ protein which manifested mostly spherical globules with smooth margins (**Figure 8A** and **Supplementary figure 13S**). Notably, SO_4_^2-^ ions could also enhance the full-length TDP-43’s aggregation levels when incubated with 120 mM SO_4_^2-^ ions and the sizes of the aggregates were found furthermore increased upon increasing the SO_4_^2-^ ion concentration to 200 mM (**Figure 8A** and **Supplementary figure 13S**).

**Figure 8.**
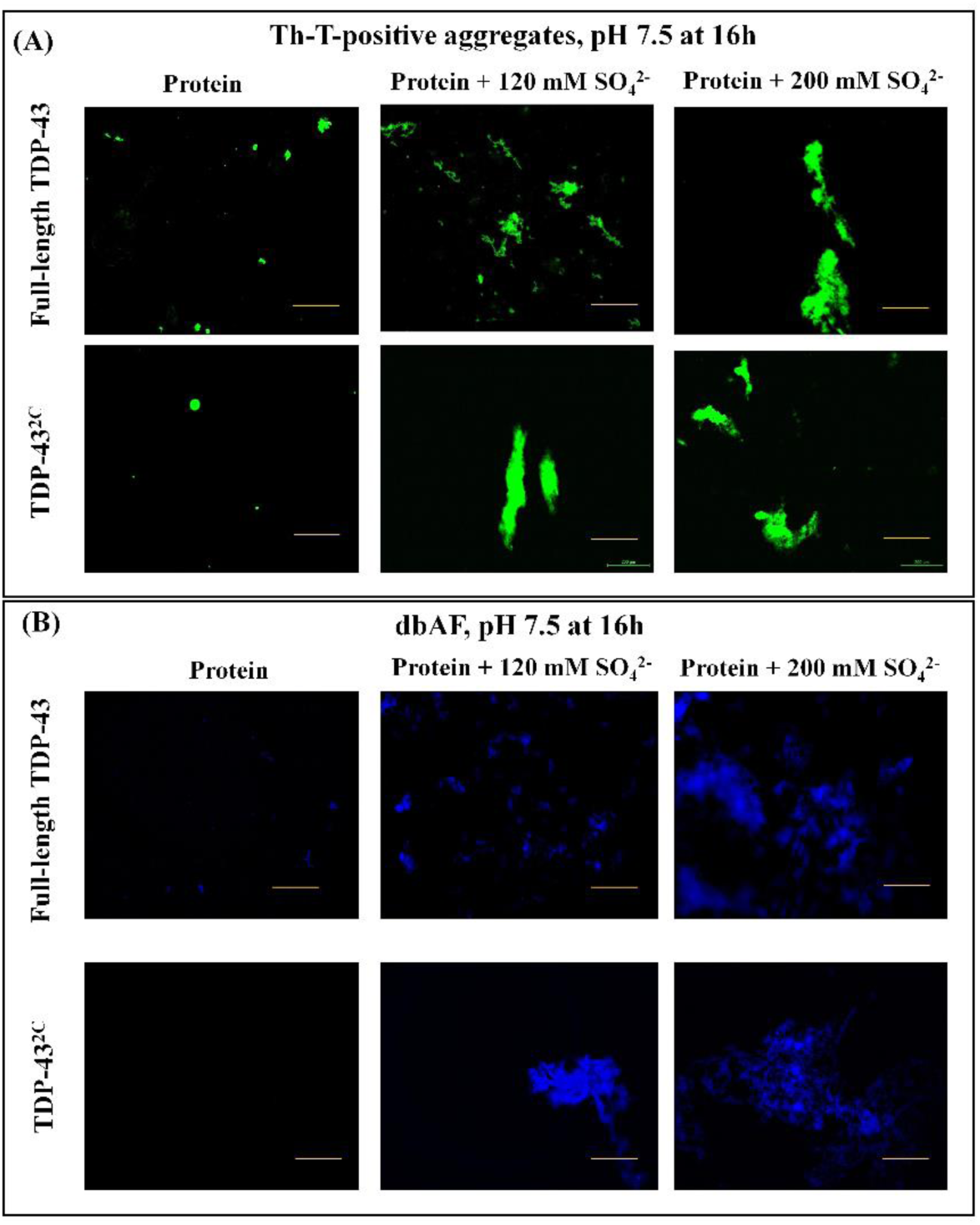
Visualization of thioflavin-T-positive full-length TDP-43 and TDP-43^2C^ aggregates and their deep-blue autofluorescence (dbAF) upon incubation with SO_4_^2-^ ions at pH 7.5. **(A)** Visualization of the Th-T-positive green fluorescent aggregates of full-length TDP-43 (50 µM) and TDP-43^2C^ ((400 µM) obtained in absence or presence of SO_4_^2-^ ions (120 mM and 200 mM) after 16h of incubation in the aggregation buffer pH 7.5. For recording the fluorescence images, the GFP filter of Leica DM2500 fluorescence microscope and 10x objective lens were used followed by color processing and background subtraction of the images using ImageJ. Scale bar is 200 µm. **(B)** Fluorescence micrographs of deep-blue autofluorescence (dbAF)-positive fluorescent aggregates of full-length TDP-43 (50 µM) and TDP-43^2C^ ((400 µM) obtained in absence or presence of SO_4_^2-^ ions (120 mM and 200 mM) after 16h of incubation in the aggregation buffer pH 7.5. For recording the fluorescence images, the UV filter of Leica DM2500 fluorescence microscope, 10x objective lens and imaging exposure time of 3s, were used followed by color processing of the images using ImageJ. Scale bar is 200 µm.

In view of the enhanced aggregations of both the full-length TDP-43 as well as TDP-43^2C^ observed in presence of SO_4_^2-^ ions, when we examined these aggregates for dbAF using the UV filter in fluorescence microscopy, the full-length TDP-43 samples incubated with 120 mM SO_4_^2-^ ions manifested relatively larger size and a greater number of aggregates emitting dbAF compared to the samples lacking SO_4_^2-^ ions during the incubation for aggregation **(Figure 8B** and **Supplementary figure 14S)**. The sizes of the full-length TDP-43 aggregates emitting dbAF were furthermore increased when the level of the SO_4_^2-^ ions was increased to 200 mM **(Figure 8B** and **Supplementary figure 14S)**. Similarly, incubation with 120 mM as well as 200 mM SO_4_^2-^ ions caused dramatic enhancement of the dbAF emission by TDP-43^2C^ which, in contrast, when incubated without the addition of any SO_4_^2-^ ions did not manifest any dbAF-positive aggregates (**Figure 8B** and **Supplementary figure 14S**). Taken together, the data support that the enhancement of the dbAF of TDP-43 or its CTF can happen whenever the aggregation is enhanced irrespective of the enhancer being a metal ion, such as Zn^2+^, or a non-metal ion such as SO_4_^2-^.

## 4. Conclusion

In this study, we examined if the *in vitro* enhancement of the TDP-43 aggregation can generate or enhance intrinsic deep-blue autofluorescence (dbAF) previously reported for a few other protein aggregates and whether dbAF is emitted by all or only liquid-like or solid-like TDP-43 assemblies or aggregates of TDP-43. We first tested the *in vitro* enhancement of the aggregations of full-length TDP-43 and its two C-terminal fragments (CTFs), TDP-43^2C^ (aa: 193-414) and the TDP-43-low complexity domain (LCD) (aa: 274-414) utilizing a gamut of tools such as thioflavin-T (Th-T) fluorescence, turbidimetry, atomic force microscopy and fluorescence microscopy of Alexa Fluor-labelled protein. In view that Zn^2+^ was previously documented to enhance the *in vitro* aggregations of certain domains of TDP-43 and several metal ions, including Zn^2+^ and Mn^2+^, are also linked to the metal dyshomeostasis in TDP-43 proteinopathies like ALS, we tested the enhancement of *in vitro* aggregation and any concurrent dbAF generation by full-length TDP-43 and its CTFs in presence of Zn^2+^ and Mn^2+^. We found enhancement in the *in vitro* solid-like aggregations of the full-length TDP-43 and TDP-43^2C^ in presence of Zn^2+^ and Mn^2+^ which also concurrently enhanced the emission of dbAF. On the contrary, Alexa fluor-633-labeled-TDP-43-LCD manifested a quick phase separation into droplets in the presence of Zn^2+^ ions and these instantaneously phase-separated species failed to emit dbAF. However, when Alexa fluor-633-labeled-TDP-43-LCD phase-separated assemblies, upon further incubation, matured into solid-like but non-amyloid nature aggregates they also emitted dbAF. Likewise, the presence of a kosmotropic ion, SO_4_^2-^, which was previously reported to enhance the *in vitro* aggregation of TDP-43^2C^, also enhanced the aggregation of full-length TDP-43 and the SO_4_^2-^-induced enhanced aggregations also concomitantly enhanced the dbAF generation by the full-length TDP-43 as well as TDP-43^2C^ proteins thereby ruling out the enhancement of dbAF of these proteins observed in presence of metal ions being due to the mere presence of the Zn^2+^ and Mn^2+^ ions. Atomic force microscopy of TDP-43^2C^ samples incubated with or without Zn^2+^ and Mn^2+^ ions also suggested presence of only small oligomers at 1h of incubation in the samples lacking the metal ions whereas those incubated for similar duration with Zn^2+^ and Mn^2+^ revealed the presence of larger-size particles thereby suggesting that oligomers fail to emit dbAF whereas larger aggregates are capable of emitting dbAF. Notably, the TDP-43 aggregates of both amyloid and non-amyloid nature, but not the oligomers or the phase-separated droplets of TDP-43, manifested dbAF. The observed enhancement of dbAF property of full-length TDP-43 and its CTFs in the presence of *in vitro* enhancers could be utilized as a label-free aggregation detection tool. Also, enhanced dbAF property could also be potentially employed to differentiate between non-pathogenic, liquid-like physiologically relevant protein assemblies and pathogenic solid-like aggregates of the full-length TDP-43 and its CTF proteins.

## Acknowledgments

We thank IIT Hyderabad, funded by Ministry of Education (MoE), Govt. of India, for research infrastructure and support. PS thanks MoE, Govt. of India, for senior research fellowship (SRF). PV thanks MoE, Govt. of India for research assistantship. VB thanks DBT, Govt. of India for SRF. We thank Yoshiaki Furukawa, Keio University, Japan, for plasmid gift. We thank S. Chengappa Thumisi, IIT-Hyderabad, for help with AFM imaging. Basant K Patel thanks SERB-DST, Govt. of India for a research grant (Grant no: SERB/CRG/2021/006856).

