## Supplemental figures for "Enhanced *in vitro* aggregation, but not phase separation, of TDP-43 and its C-terminal fragments generate deep-blue autofluorescence"

**Title**

Basant K Patel, PhD

Professor of Biotechnology

### Supplementary figure 1S

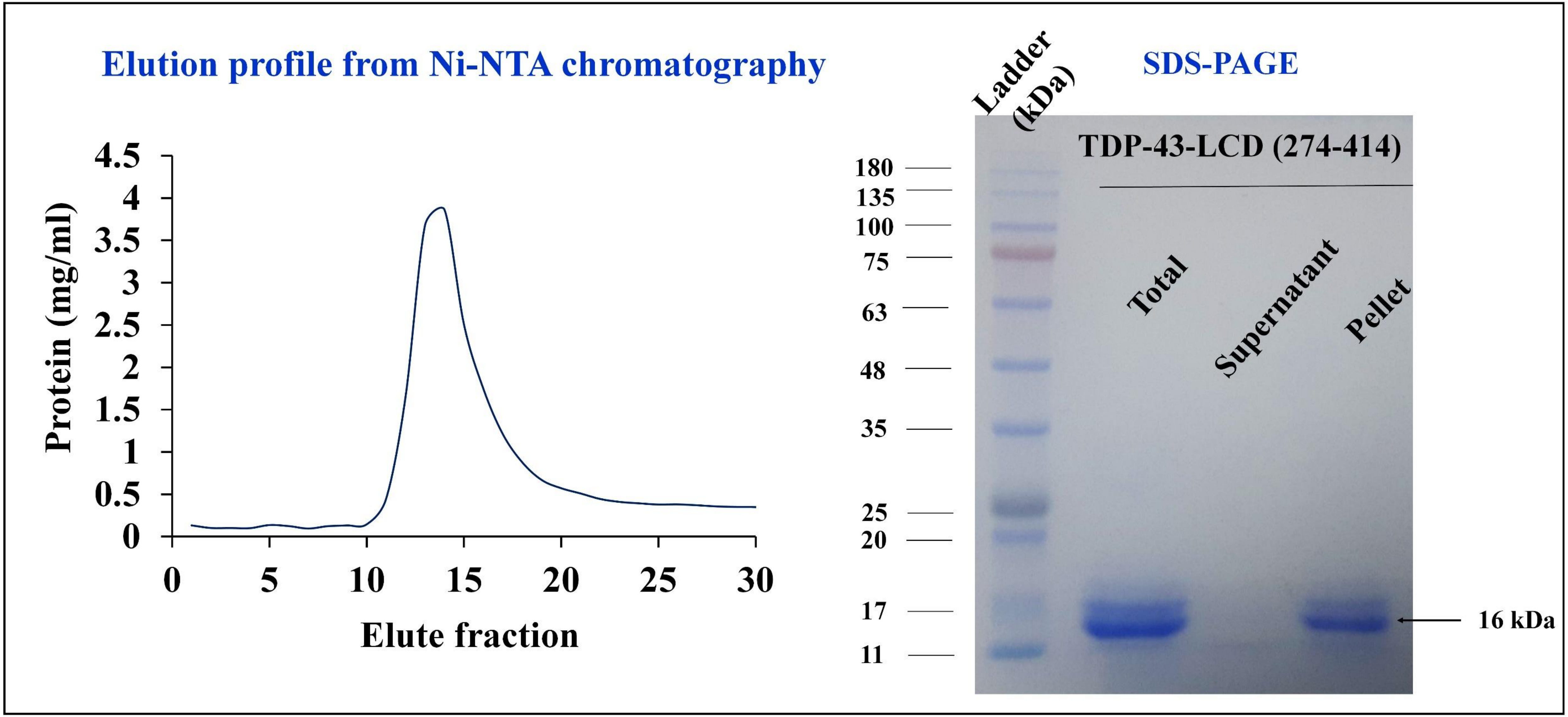

**Supplementary figure 1S. Recombinant expression and purification of TDP-43-LCD using Nickel affinity chromatography.**

**Left panel-** Detection of thee TDP-43-LCD protein levels in the eluted fractions from Ni-NTA chromatography. The protein concentration (mg/mL) was obtained from the absorbance at 280 nm by using the molar extinction co-efficient of 17990 $M^{-1}cm^{-1}$. **Right panel-** The eluted fraction with the highest protein concentration was dialyzed against PBS for 24h where the protein was observed to precipitate. The precipitated sample was centrifuged at 20,000 x g for 30 minutes and the pellet and the supernatant fractions were separately collected and analyzed for protein homogeneity by 12% SDS-PAGE along side molecular weight standards.

#### Supplementary figure 2S

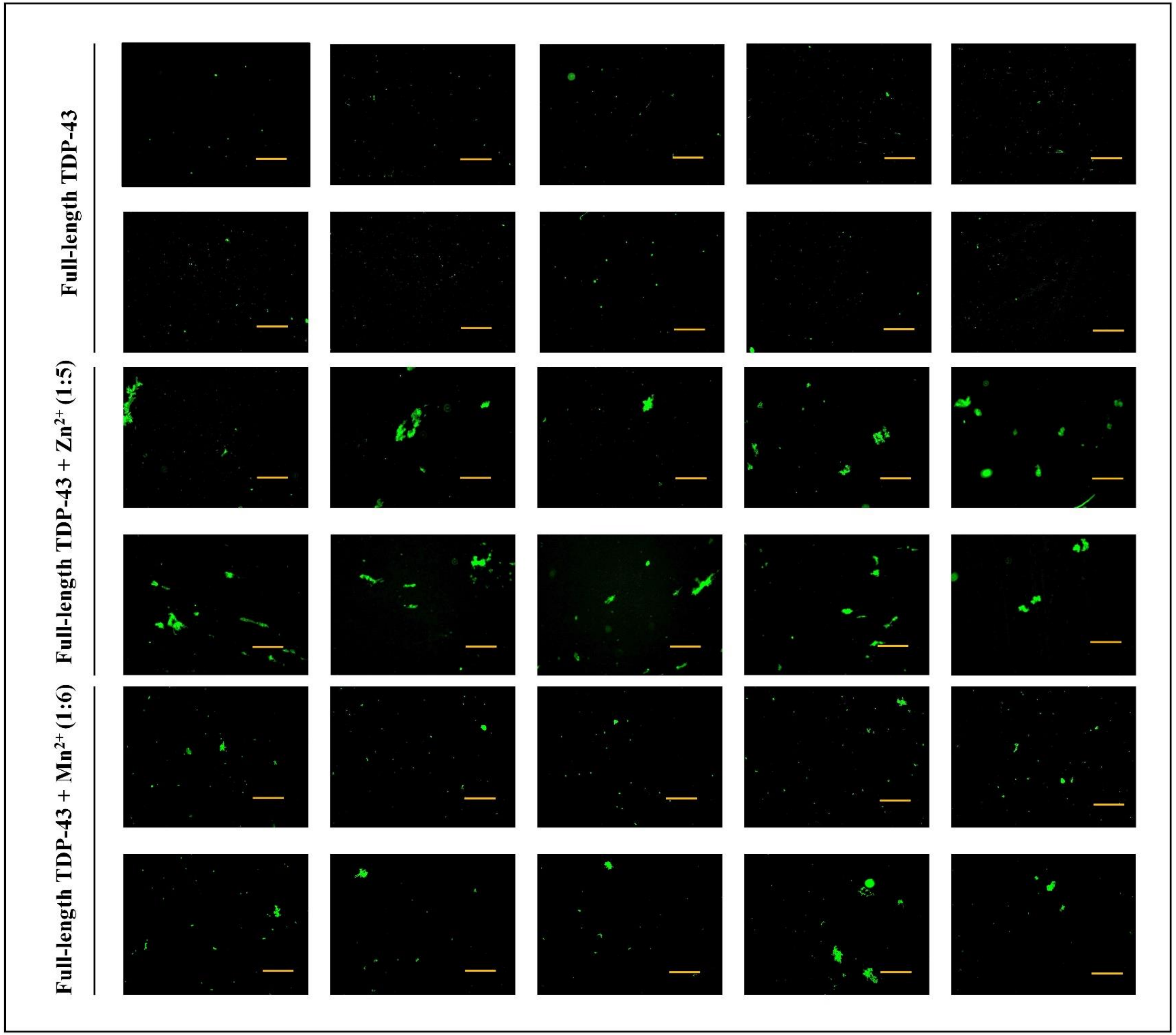


**Supplementary figure 2S. Thioflavin-T (Th-T)-positive aggregates of full-length TDP-43 in the presence of $Zn^{2+}$ and $Mn^{2+}$ at pH 7.5.**

After 16h incubation, the aggregated samples of full-length TDP-43 in the absence or presence of $Zn^{2+}$ and $Mn^{2+}$, were examined for the presence of Th-T-positive green fluorescent aggregates under the GFP filter in the Leica DM2500 fluorescence microscope. The images were acquired using the 10x objective lens. Scale bar – 200 µm. All images were processed using ImageJ.

Supplementary figure 3S

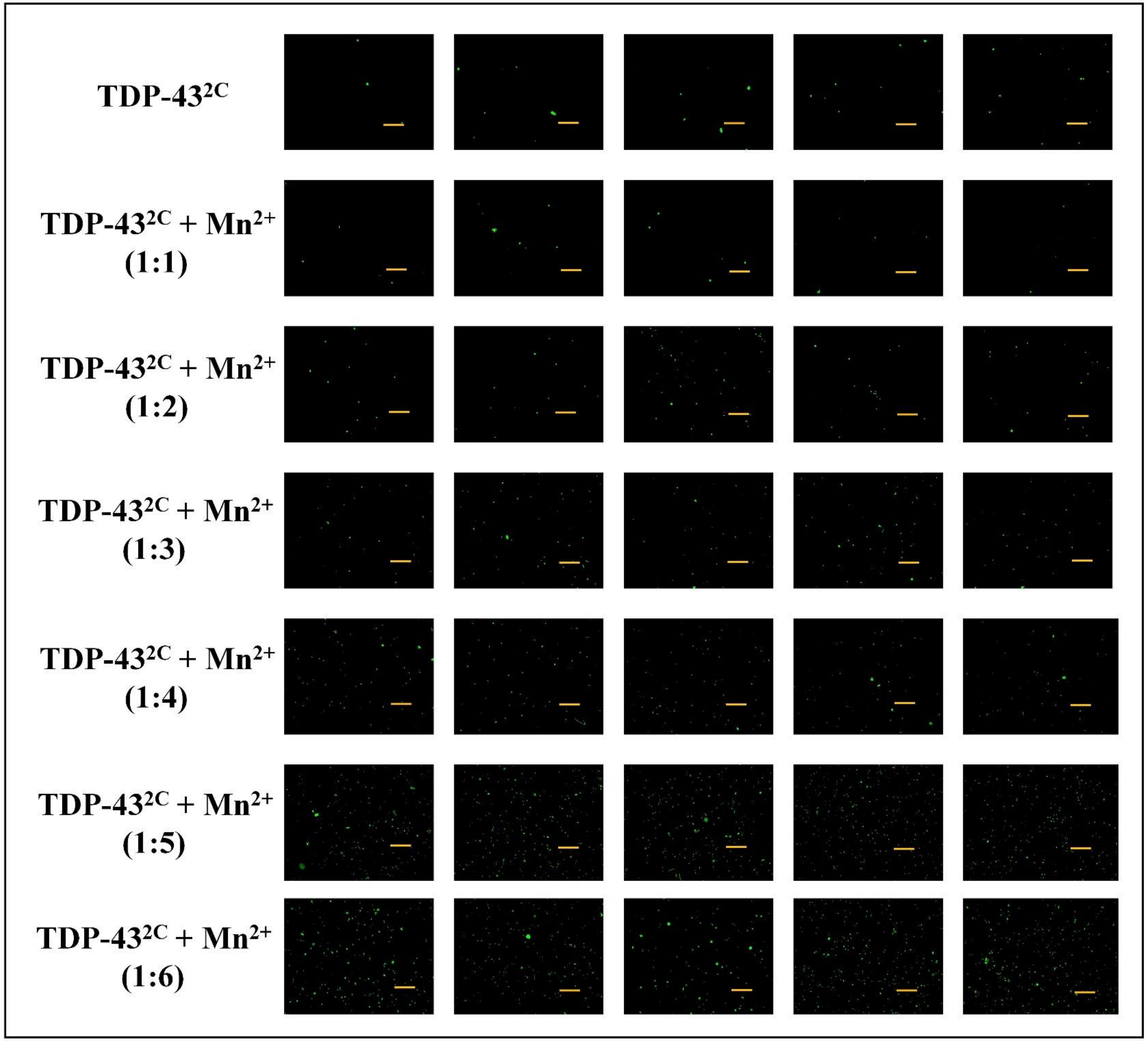


**Supplementary figure 3S. Thioflavin-T (Th-T)-positive aggregates of TDP-43$^{2C}$ in the presence of $Mn^{2+}$ at pH 7.5.**

After 16h incubation, aggregated samples of TDP-43$^{2C}$ in absence or presence of different stoichiometric ratios of $Mn^{2+}$ (1:1-1:6), were examined for the presence of Th-T-positive green fluorescent aggregates under the GFP filter in the Leica DM2500 fluorescence microscope. The images were acquired using the 10x objective lens. Scale bar – 200 µm. All images were processed using ImageJ.

#### Supplementary figure 4S

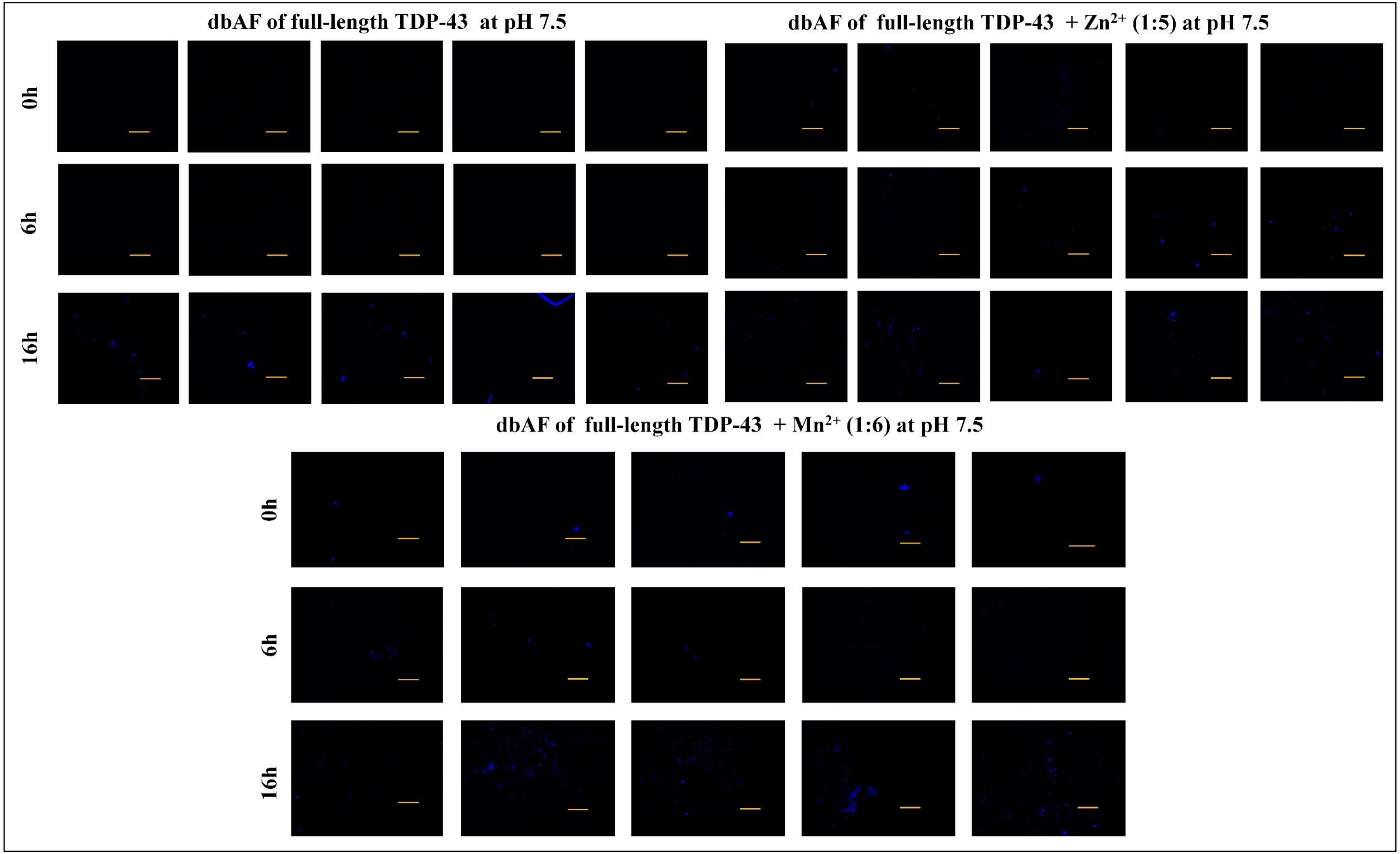

**Supplementary figure 4S. Deep-blue autofluorescence (dbAF) of full-length TDP-43 in the presence of $Zn^{2+}$ and $Mn^{2+}$ at pH 7.5.** The aggregated samples of full-length TDP-43 obtained with or without $Zn^{2+}$ and $Mn^{2+}$ addition, were examined for the presence of aggregated structures manifesting dbAF using the UV filter in the Leica DM2500 fluorescence microscope. The samples were imaged at different time intervals of incubation such as 0h, 6h and 16h. The images were acquired using the 10x objective lens. Scale bar – 200 µm. All the images were processed using ImageJ.

#### Supplementary figure 5S

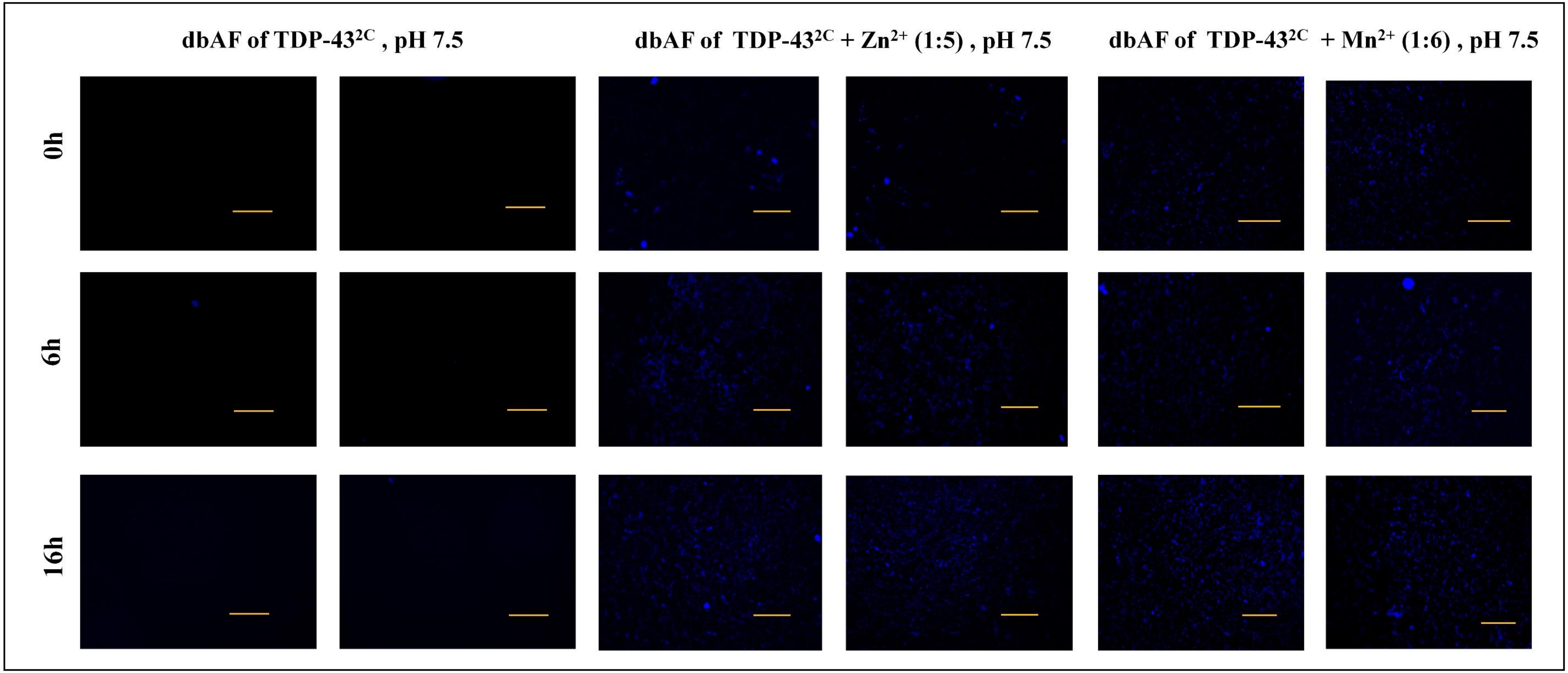

**Supplementary figure 5S. Deep-blue autofluorescence (dbAF) of TDP-43$^{2C}$ in the presence of $Zn^{2+}$ and $Mn^{2+}$ at pH 7.5.**

The aggregated samples of TDP-43$^{2C}$ with or without the incubation with $Zn^{2+}$ and $Mn^{2+}$ , were examined for the presence of aggregated structures manifesting dbAF using the UV filter in Leica DM2500 fluorescence microscope. The samples were imaged at different time intervals of incubation such as 0h, 6h and 16h. The images were acquired using the 10x objective lens. Scale bar – 200 µm. All the images were processed using ImageJ.

**Supplementary figure 6S**

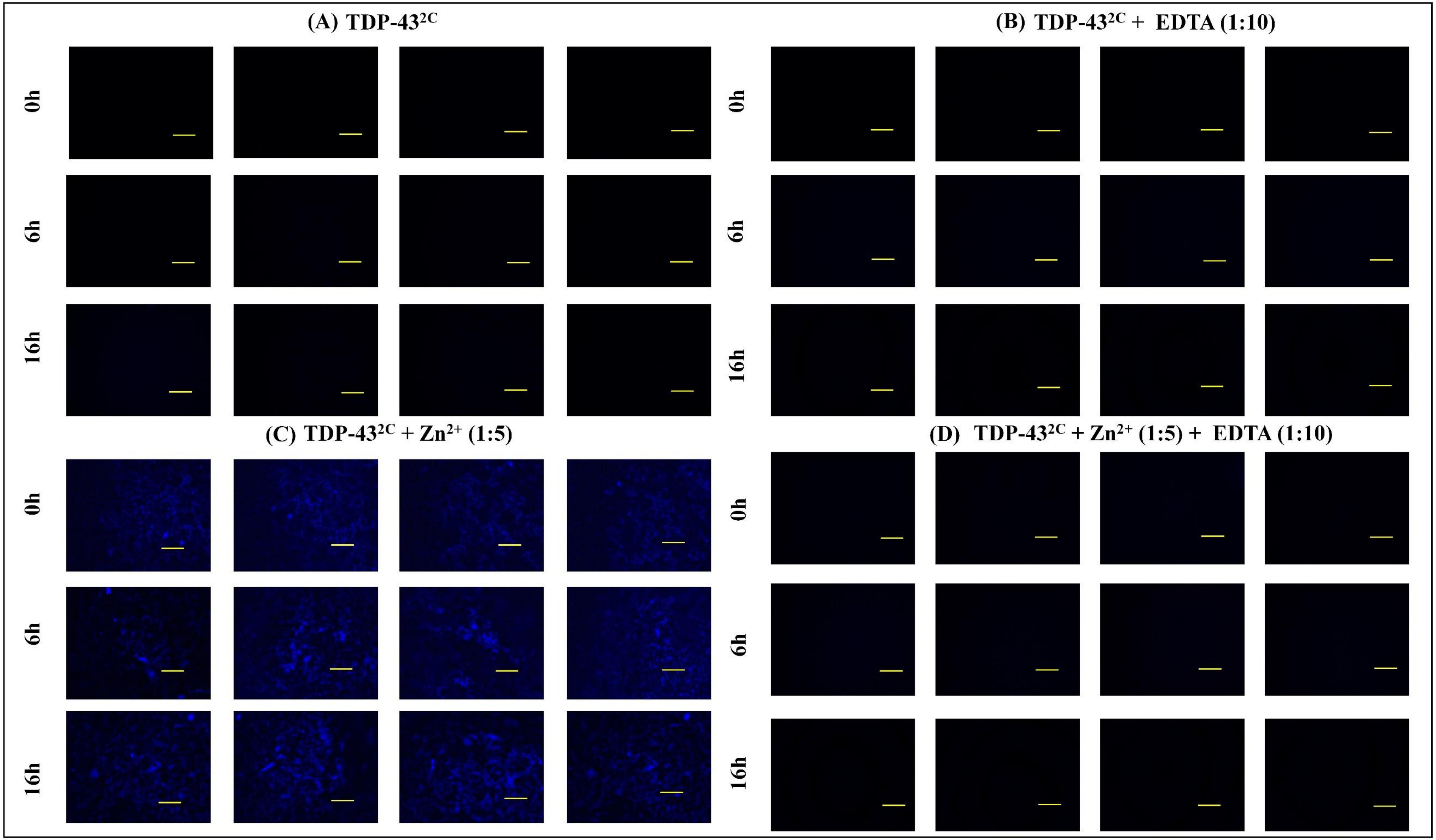

**Supplementary figure 6S. Deep-blue autofluorescence (dbAF) of TDP-43$^{2C}$ in presence of metal chelator, EDTA, and $Zn^{2+}$ at pH 7.5.** The samples of TDP-43$^{2C}$ were incubated either in absence or presence and of $Zn^{2+}$ (protein: $Zn^{2+}$ = 1:5) and both types of these samples were also added with a metal chelator, EDTA, (protein: EDTA = 1:10) and then visualized at 0h, 6h and 16h of incubation for dbAF emission under the UV filter of the Leica DM2500 fluorescence microscope. The images were acquired using the 10x objective lens. Scale bar – 200 µm. All the images were background subtracted and processed for color using ImageJ.

### Supplementary figure 7S

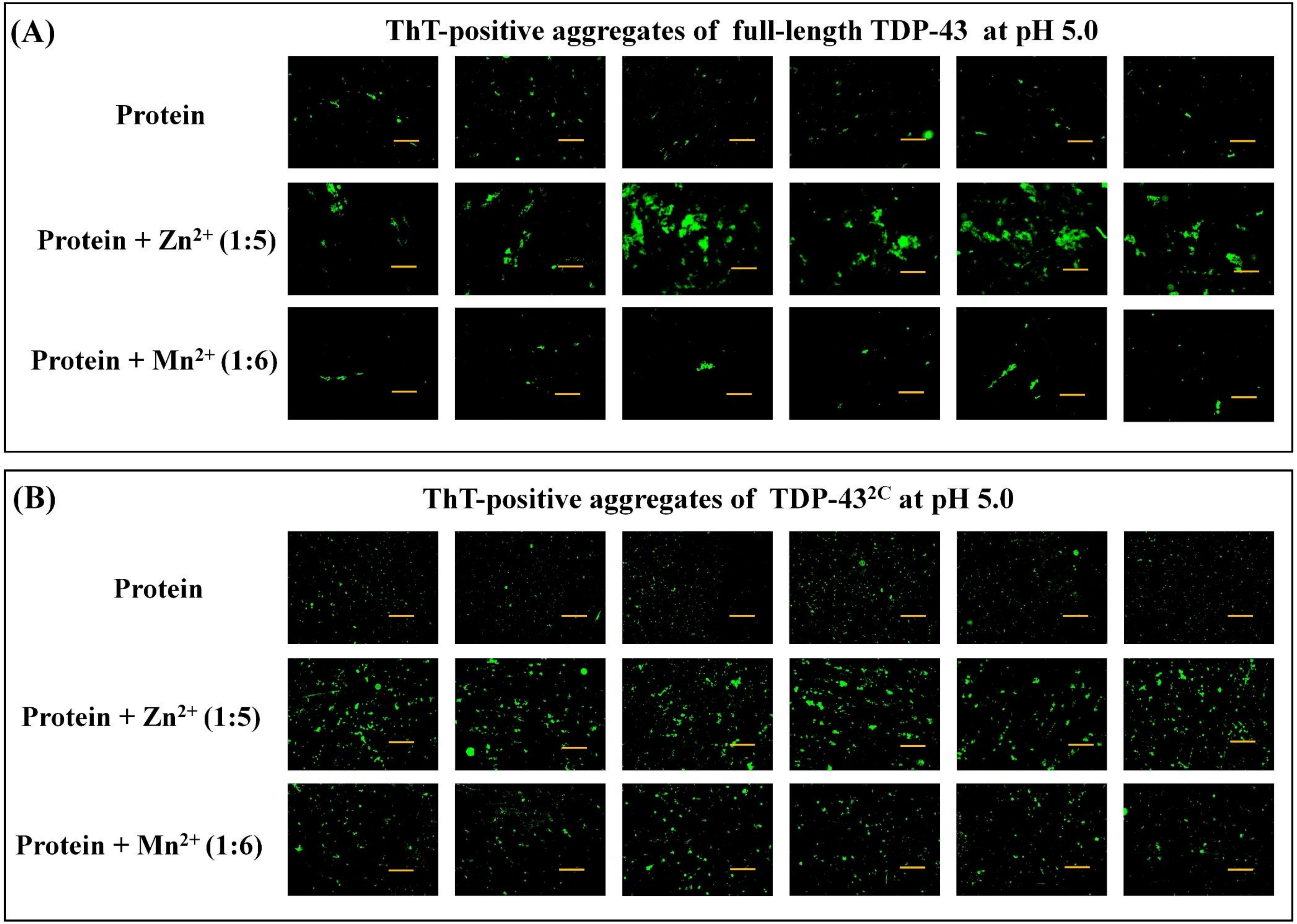

**Supplementary figure 7S. Thioflavin-T-positive aggregates of full-length TDP-43 and TDP-43$^{2C}$ in the presence of $Zn^{2+}$ and $Mn^{2+}$ at pH 5.0.**

**(A)** After 16h incubation, the aggregated samples of full-length TDP-43 obtained in absence or presence of $Zn^{2+}$ and $Mn^{2+}$, were examined for the presence of Th-T-positive green fluorescent aggregates under the GFP filter in Leica DM2500 fluorescence microscope. The images were acquired using the 10x objective lens. Scale bar – 200 µm. All images were processed using ImageJ. **(B)** After 16h incubation, the aggregated samples of TDP-43$^{2C}$ obtained in absence or presence of $Zn^{2+}$ and $Mn^{2+}$, were examined for the presence of Th-T-positive green fluorescent aggregates under the GFP filter in Leica DM2500 fluorescence microscope. The images were acquired using the 10x objective lens. Scale bar – 200 µm. All images were processed using ImageJ.

#### Supplementary figure 8S

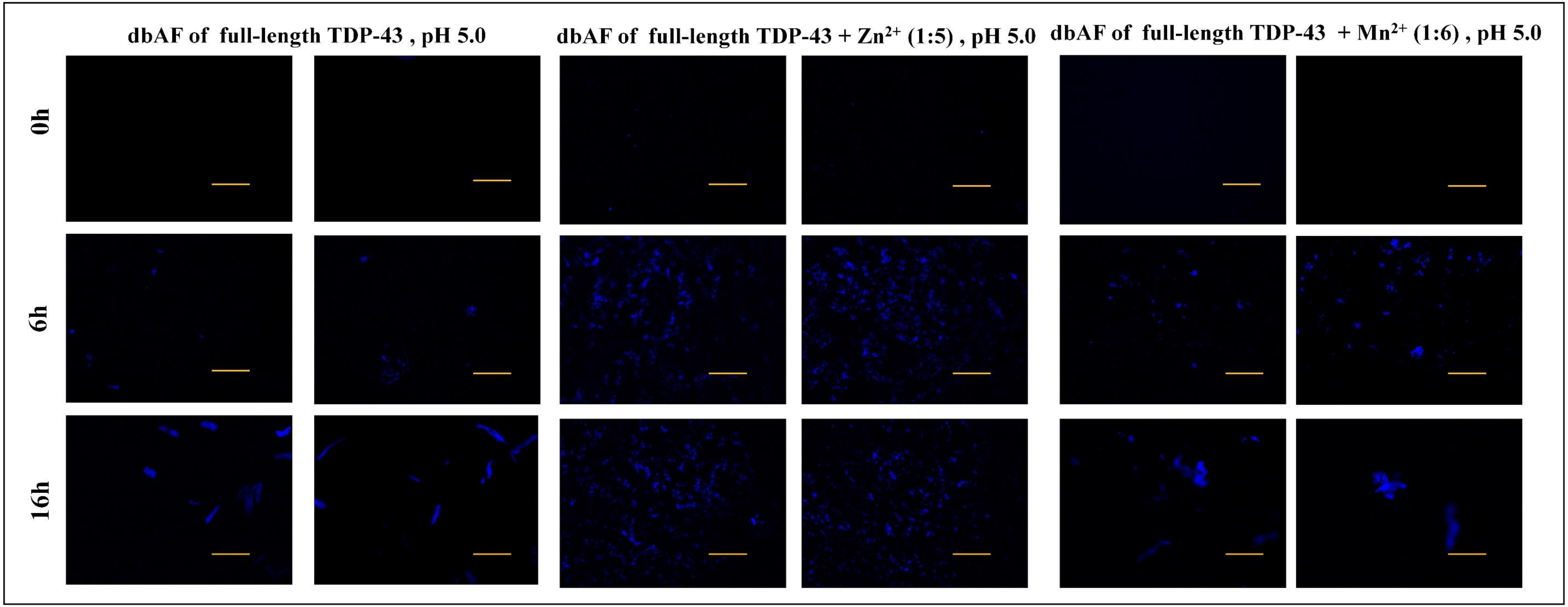

**Supplementary figure 8S. Deep-blue autofluorescence (dbAF) of full-length TDP-43 in the presence of $Zn^{2+}$ and $Mn^{2+}$ at pH 5.0.** The samples of TDP-43 aggregated with or without $Zn^{2+}$ and $Mn^{2+}$ were examined for the presence of aggregated structures manifesting dbAF using the UV filter in Leica DM2500 fluorescence microscope. The samples were collected at different time intervals such as 0h, 6h and 16h. The images were acquired using the 10x objective lens. Scale bar – 200 µm. All the images were processed using ImageJ.

#### Supplementary figure 9S

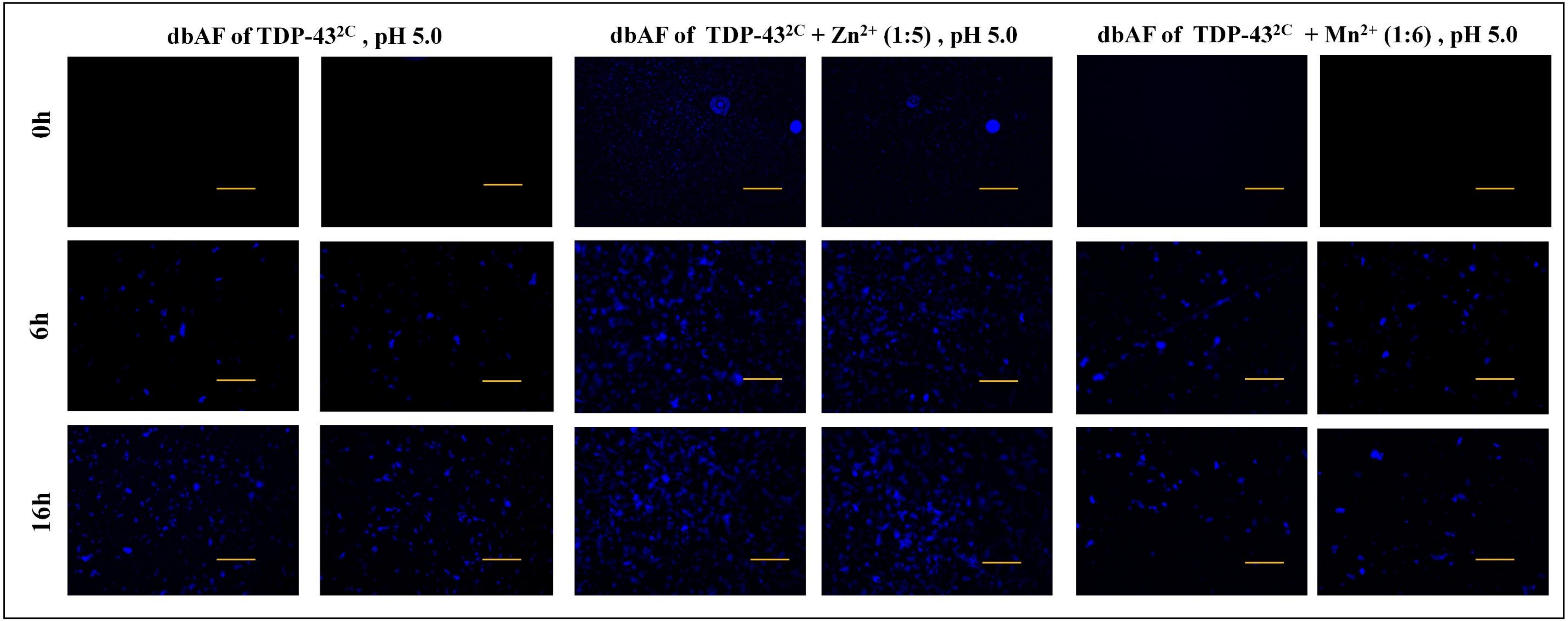

**Supplementary figure 9S. Deep-blue autofluorescence (dbAF) of TDP-43$^{2C}$ in the presence of divalent cations at pH 5.0.** The aggregated samples of TDP-43$^{2C}$ with and without $Zn^{2+}$ and $Mn^{2+}$ are examined for the presence of aggregated structures manifesting dbAF using the UV filter in Leica DM2500 fluorescence microscope The samples were collected at different incubation times- 0h, 6h and 16h. The images were acquired using the 10x objective lens. Scale bar – 200 µm. All the images were processed using ImageJ.

**Supplementary figure 10S**

#### Atomic Force Microscopy

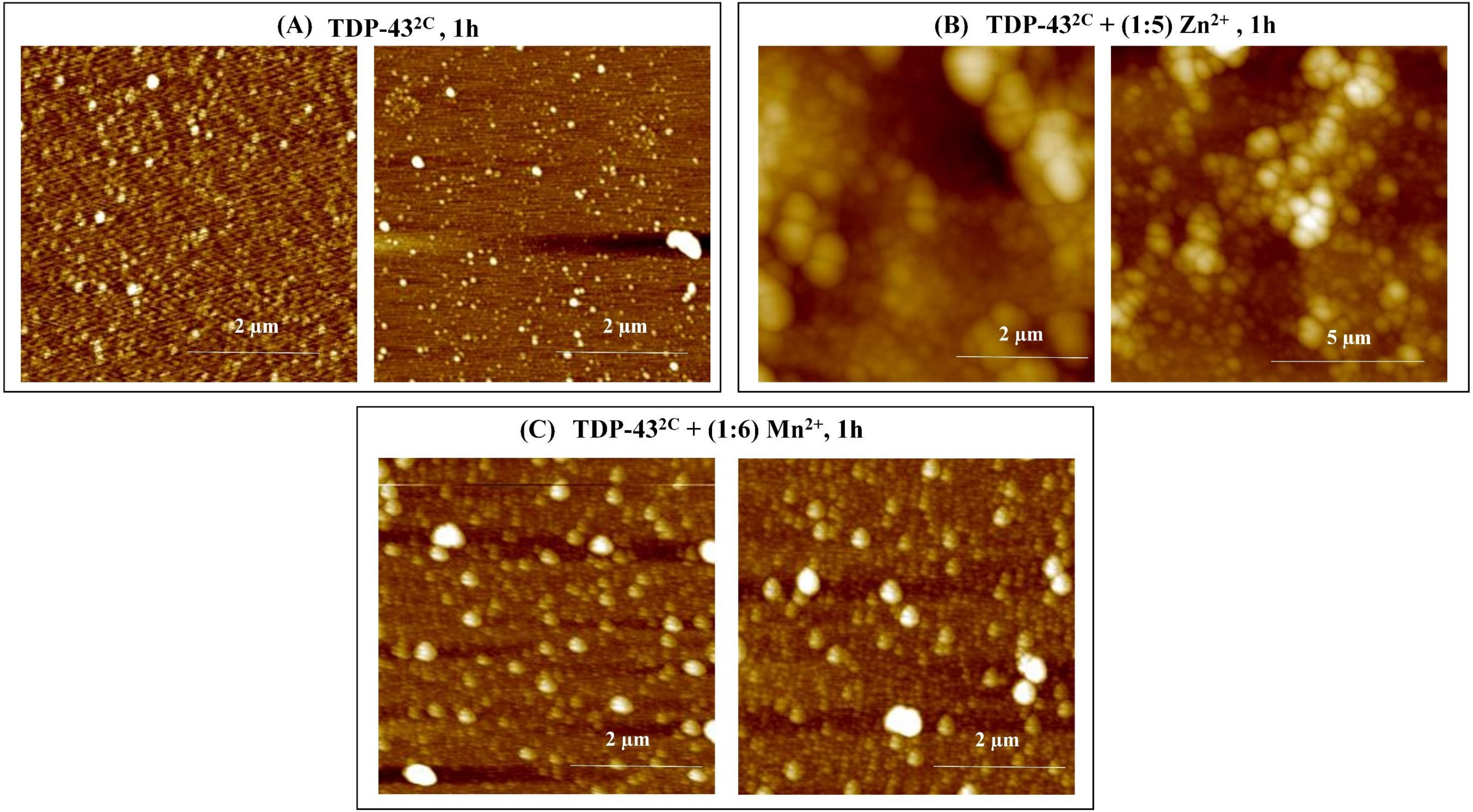

**Supplementary figure 10S. AFM imaging of TDP-43$^{2C}$ in absence or presence of $Zn^{2+}$ and $Mn^{2+}$ ions at pH 7.5.** The morphological characteristics of samples of TDP-43$^{2C}$ incubated in absence or presence and of $Zn^{2+}$ and $Mn^{2+}$ were analyzed by AFM imaging. For this, 400 µM of TDP-43$^{2C}$ was incubated in absence or presence and of $Zn^{2+}$ and $Mn^{2+}$ at the stoichiometric ratio of 1:5 (protein: $Zn^{2+}$) and 1:6 (protein: $Mn^{2+}$) respectively for one hour at 37°C with rotation at 200 rpm. **(A)** TDP-43$^{2C}$ incubated in absence of the $Zn^{2+}$ and $Mn^{2+}$ ions. Scale bar – 2 µm. **(B)** TDP-43$^{2C}$ incubated in presence of the $Zn^{2+}$ ion. Scale bar – 5 µm. **(C)** TDP-43$^{2C}$ incubated in presence of the $Mn^{2+}$ ion. Scale bar – 2 µm.

#### Supplementary figure 11S

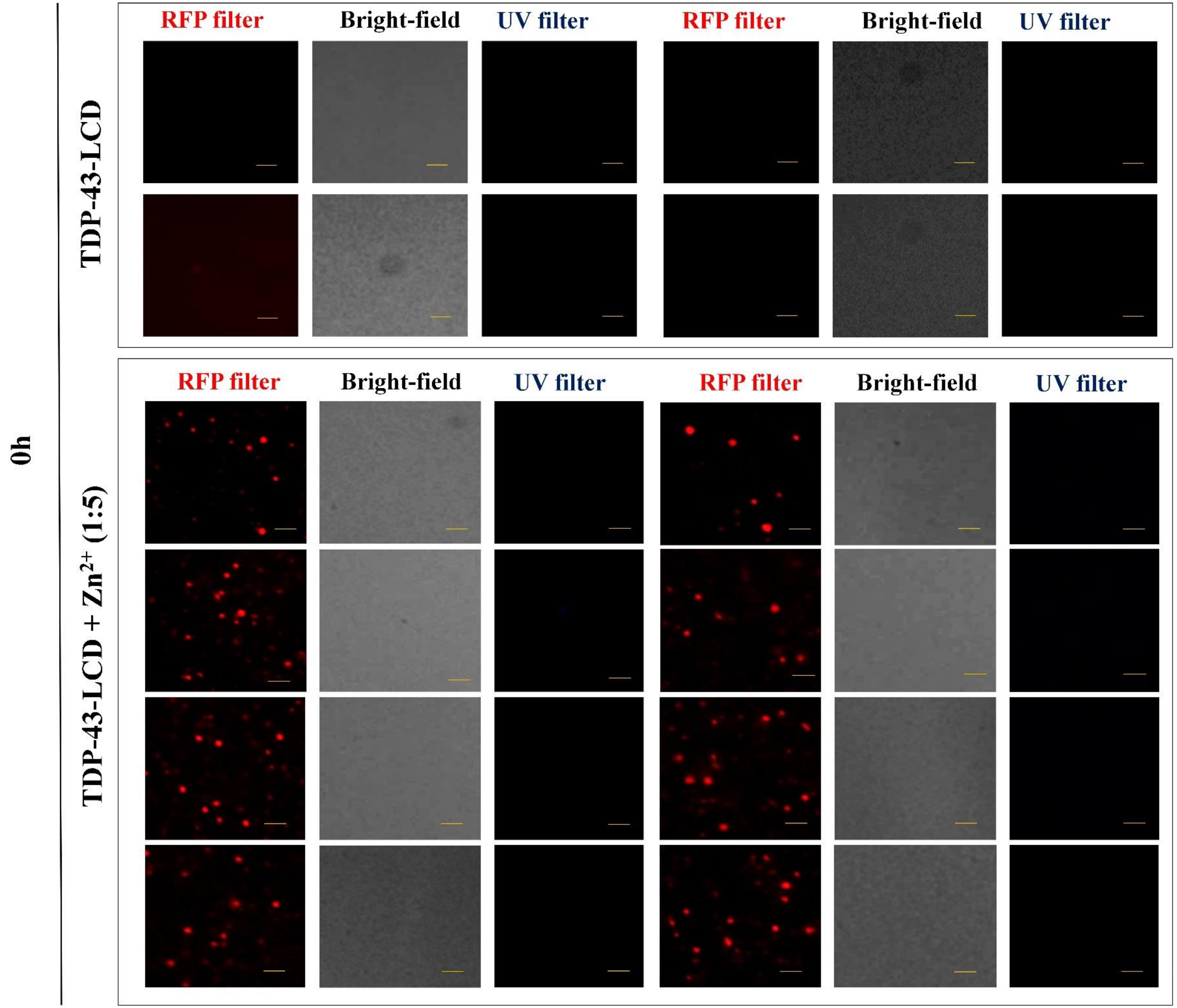

**Supplementary Figure 11S. Effect of $Zn^{2+}$ at zero hour of incubation on the *in vitro* aggregation, phase-separation and deep-blue autofluorescence (dbAF) of TDP-43-LCD.**

TDP-43-LCD was labelled with Alexa Fluor™ 633 NHS ester dye. The labelled and unlabelled proteins were mixed in the ratio of 1:15 (labelled: unlabelled) to observe any phase-separation in absence or presence and of the $Zn^{2+}$ ions (protein : $Zn^{2+}$ = 1 : 5). TDP-43-LCD (15 µM total protein, 1.5M urea, 50 mM phosphate buffer, pH 7.5) at the 0h incubation without or with $Zn^{2+}$ was visualized under RFP filter and bright-field of Leica DM2500 fluorescence microscope to detect for phase separation. Furthermore, dbAF from the aggregates/phase-separated structures were also visualized under UV filter. All the images were acquired under 20 x objective lens. Scale bar – 10 µm.

**Supplementary figure 12S**

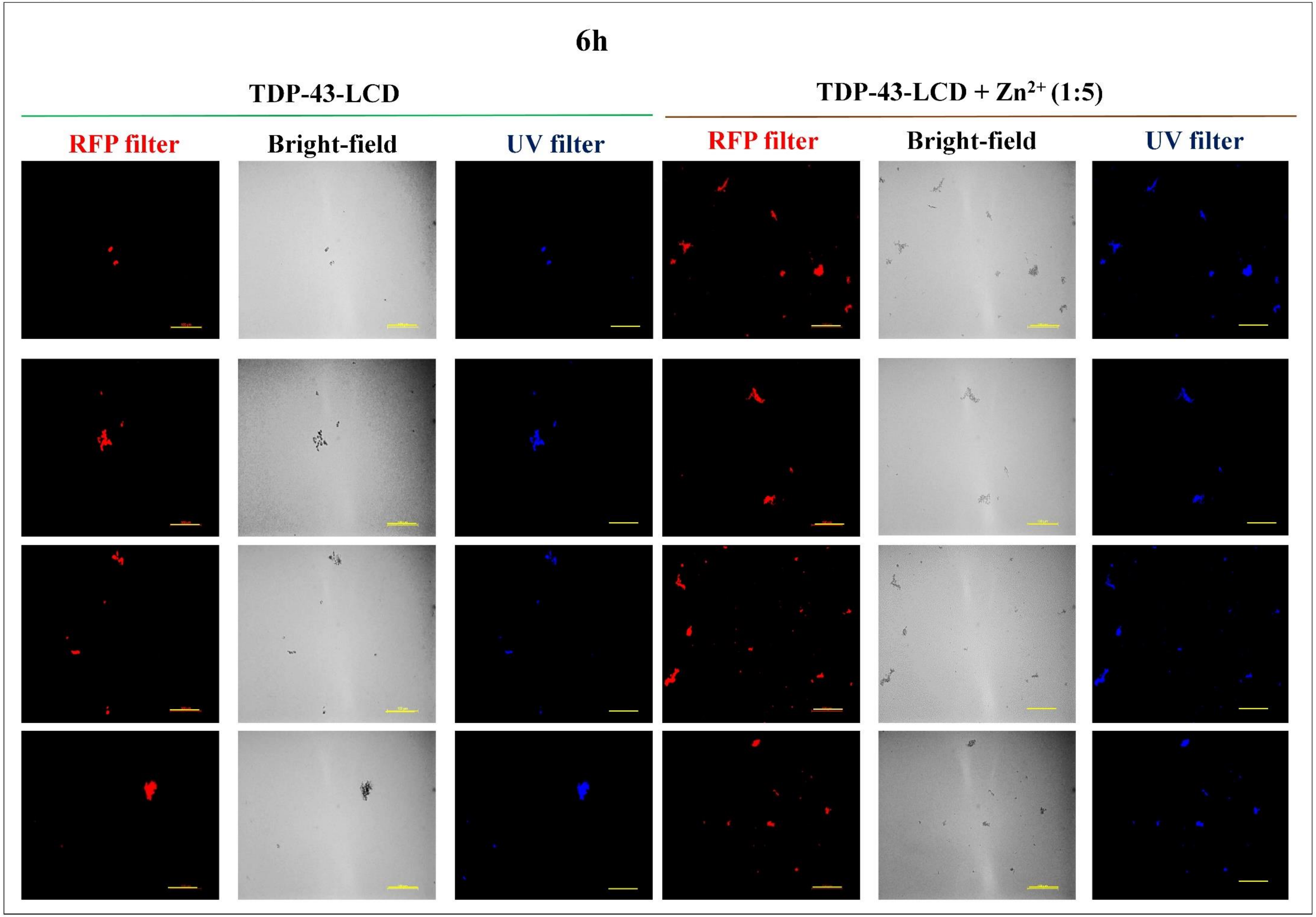

**Supplementary Figure 12S. Effect of $Zn^{2+}$ ion at 6 hour of incubation on the *in vitro* aggregation, phase-separation and deep-blue autofluorescence (dbAF) of TDP-LCD.** TDP-43-LCD was labelled with Alexa Fluor™ 633 NHS ester dye. The labelled and unlabelled proteins were mixed in the ratio of 1:15 (labelled: unlabelled) to observe any phase-separation in absence or presence and of the $Zn^{2+}$ ions (protein : $Zn^{2+}$ = 1 : 5). TDP-43-LCD (15 µM total protein, 1.5M urea, 50 mM phosphate buffer, pH 7.5) at the 6h incubation without or with $Zn^{2+}$ was visualized under RFP filter and bright-field of Leica DM2500 fluorescence microscope to detect for phase separation. Furthermore, dbAF from the aggregates/phase-separated structures were also visualized under UV filter. All the images were acquired under 20 x objective lens. Scale bar – 100 µm.

### Supplementary figure 13S

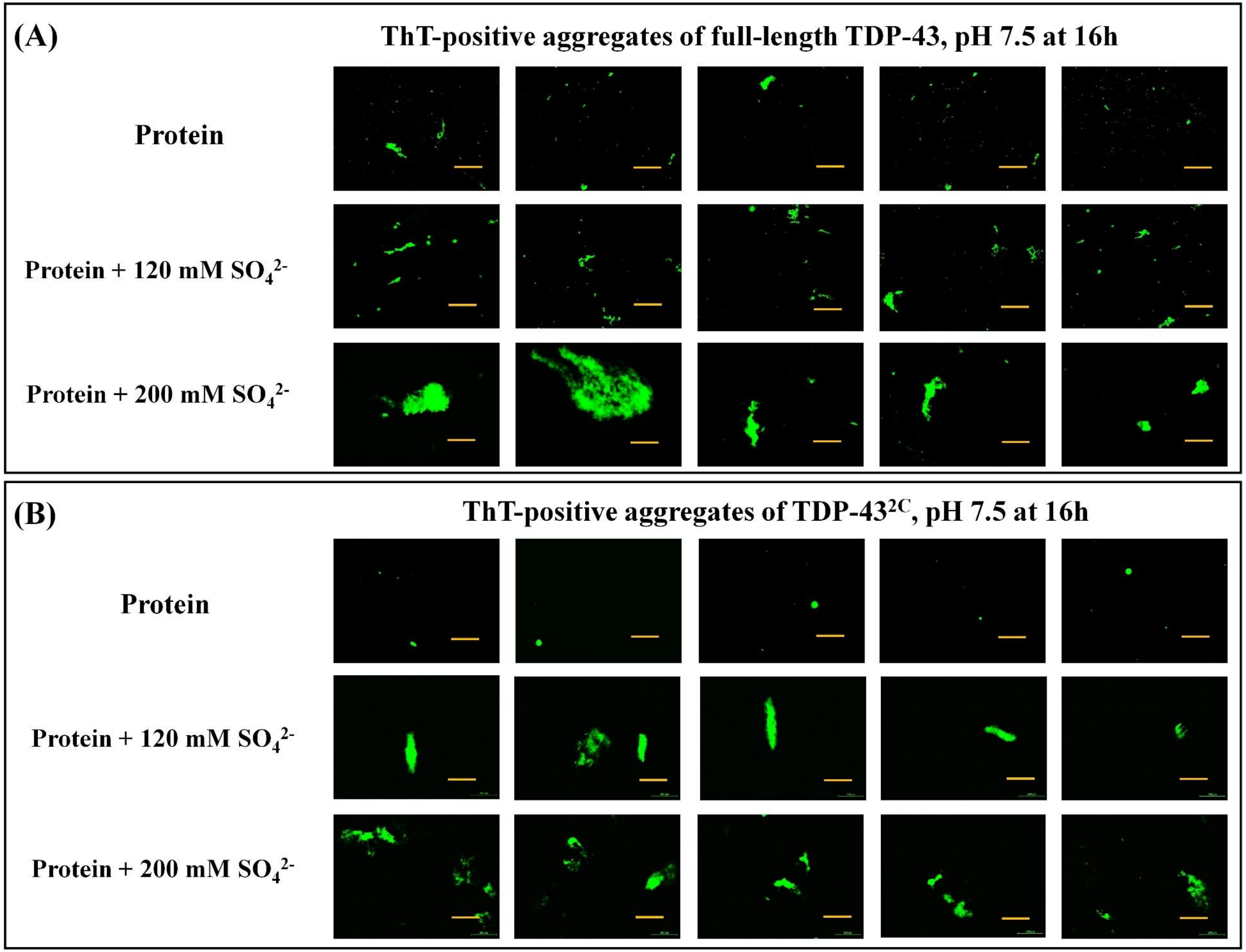

**Supplementary figure 13S. Thioflavin-T-positive aggregates of full-length TDP-43 and TDP-$43^{2C}$ in the presence of the kosmotropic anion, $SO_4^{2-}$, at pH 7.5.**

**(A)** After 16h, the samples of full-length TDP-43 aincubated in the presence of $SO_4^{2-}$ ions (120 mM and 200 mM) were examined for the presence of Th-T-positive aggregates under the GFP filter in the Leica DM2500 fluorescence microscope. The images were acquired using 10x objective lens. Scale bar – 200 µm. **(B)** After 16h, the aggregated samples of TDP-$43^{2C}$ in the presence of $SO_4^{2-}$ ions (120 mM and 200 mM) were examined for the presence of Th-T-positive aggregates under the GFP filter in the Leica DM2500 fluorescence microscope. The images were acquired using 10x objective lens. Scale bar – 200 µm.

### Supplementary figure 14S

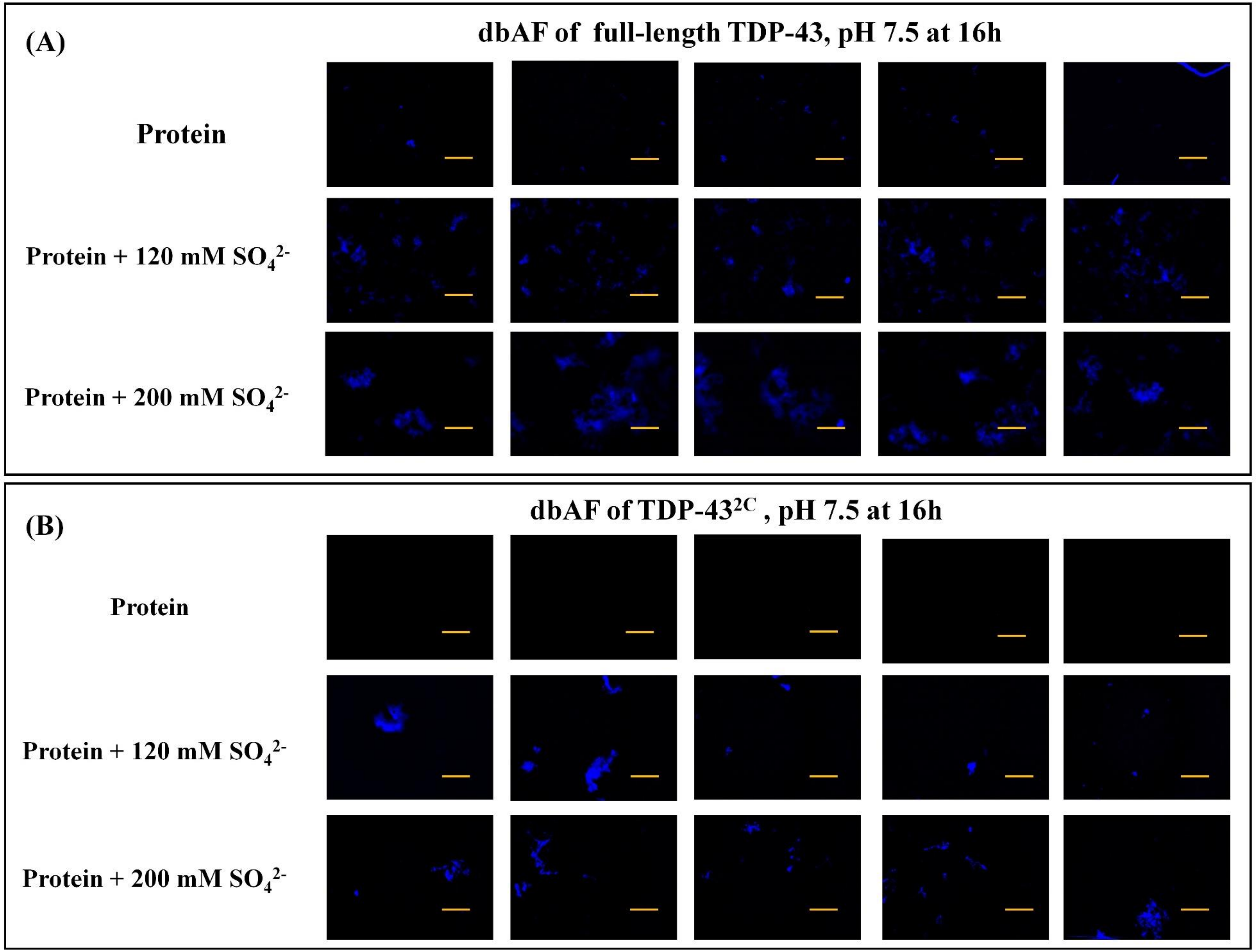

**Supplementary Figure 14S. Deep-blue autofluorescence (dbAF) of full-length TDP-43 and TDP-43$^{2C}$ in the presence of the kosmotropic anion, $SO_4^{2-}$, at pH 7.5.**

**(A)** The samples of full-length TDP-43 aggregated after 16h in absence or presence of $SO_4^{2-}$ ions (120 mM and 200 mM), were examined for the presence of dbAF emitting structures under the UV filter of the Leica DM2500 fluorescence microscope. The images were obtained under the 10x objective lens. Scale bar – 200 µm. All the images were processed using ImageJ. **(B)** The samples of TDP-43$^{2C}$ aggregated after 16h in absence or presence of $SO_4^{2-}$ ions (120 mM and 200 mM) were were examined for the presence of dbAF emitting structures under the UV filter of the Leica DM2500 fluorescence microscope. The images were obtained under the 10x objective lens. Scale bar – 200 µm. All the images were processed using ImageJ.
